# What makes a banana false? How the genome of Ethiopian orphan staple *Ensete ventricosum* differs from the banana A and B sub-genomes

**DOI:** 10.64898/2026.02.21.706659

**Authors:** Sadik Muzemil, Parameswari Paul, Laura Baxter, Ana Dominguez-Ferreras, Sunil Kumar Sahu, Allen Van Deynze, Guoqiang Mai, Zerihun Yemataw, Kassahun Tesfaye, Vardis Ntoukakis, David J. Studholme, Murray Grant

## Abstract

**Background:** *Ensete ventricosum*, also known as the “tree against hunger” plays a key role in Ethiopian food security and farming systems, feeding more than 20 million people. Since domestication via clonal selection in the south-west Ethiopian highlands, today’s diverse enset landraces contribute multiple benefits including food, fibre by-product, animal bedding and cattle fodder to farmers and local communities. Improved genomic resources for this highly drought-tolerant plant are essential to supplement the conventional clonal selection-based breeding programme and pave the way towards targeted breeding.

**Results:** We sequenced the genome of enset landrace Mazia, which is partially resistant/tolerant to Xanthomonas wilt and predicted 38,940 protein-coding genes. The Mazia assembly (540.14 Mb) is more complete than the previously published genome assembly of landrace Bedadeti (451.28 Mb) and displayed 1.41% heterozygosity and 64.64% repetitive DNA content. Comparative analyses with the Bedadeti assembly and chromosome-level genome sequences of the two main banana progenitors (*Musa acuminata*, AA genome; *Musa balbisiana*, BB genome) unexpectedly revealed ∼25% of the Mazia genome is unique to enset. Gene Ontology (GO) and sequence similarity search analysis of enset-specific protein-coding genes identified distinct functional signatures that underpin the lifestyle, adaptation, and corm productive quality of enset, including functions related to DNA integration, carbohydrate metabolism, disease resistance and transcriptional regulation. In contrast, *Musa*-specific genes showed enrichment for defence response, protein phosphorylation and fruit development pathways. Focusing on the classical nucleotide binding site leucine rich repeat (NLR) disease resistance genes, we identified and characterised NLRs in enset and *Musa* species genomes, revealing a considerable expansion in the *Musa acuminata* genome. We also identified unique genes in enset and banana genomes whose functional and evolutionary roles are yet to be determined.

**Conclusions:** Here, we report a *de novo* genome assembly for the enset (*Ensete ventricosum*) landrace Mazia and provide a high-quality annotation of both Mazia and the previously published assembly of the landrace Bedadeti. Collectively, these genomic resources provide a valuable foundation for comparative genomics within the *Musaceae* family and open new opportunities for the development of marker-assisted breeding strategies to accelerate the improvement of agronomically important traits in enset.

## 1 Background

The farmers’ adage, “Enset is our food, our clothes, our beds, our houses, our cattle-feed, our plates” [18], captures the multifaceted importance of enset (*Ensete ventricosum*), an orphan crop that serves as a staple food source for over 20 million people [17] in Africa’s second most populous country. With sub-Saharan Africa’s population projected to surpass 1.9 billion by 2050 [145], understanding and improving underutilised crops like enset is critical for regional food security.

Enset, also known as false banana or Abyssinian banana, is the world’s largest herbaceous perennial monocarpic plant, reaching heights up to 11 m. A member of the Musaceae (banana family), enset is estimated to have diverged from *Musa* species between 51.9 [65] to 59.9 [39] million years ago, and represents one of the most socioeconomically and culturally important species. Unlike banana, which is grown for its fruit, domesticated enset is cultivated for its large starch-rich underground corm and overlapping leaf sheaths (pseudostem) (Figure 1). These carbohydrate-rich components provide a staple food with high nutritional value: daily consumption of 0.5 kg of enset-derived products supplies approximately 20% of protein and 70% of energy requirements [108].

**Figure 1:**
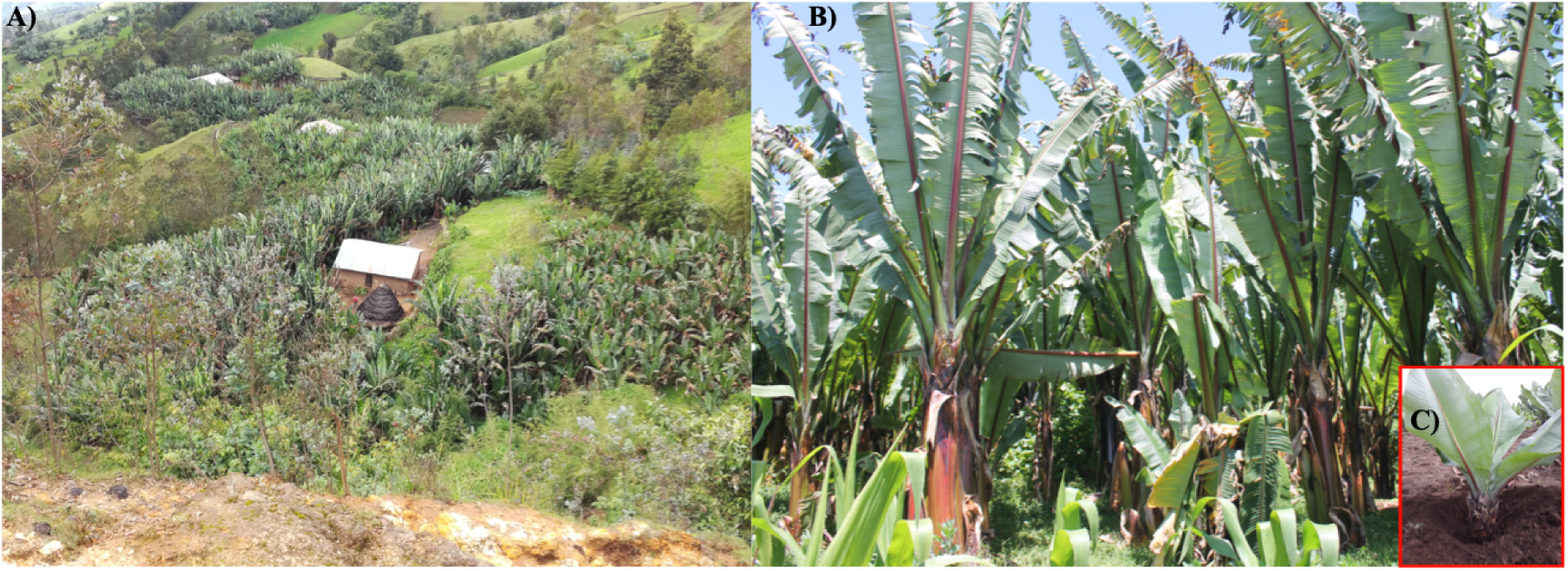
Overview of enset-based farming system in southern Ethiopia. (A) A homestead enset plantation in Dawro district containing a variety of clonally propagated landraces, each specifically selected for their preferred uses. (B) A range of clonally propagated enset plants approaching harvest growing amongst selected landraces at different developmental stages. Each landrace can be identified by its phenotypic traits such as growth habit, colour of leaf, midrib and/or leaf margins, as well as their preferred uses (e.g. harvestable plant material or tolerance to disease and drought). (C) The 2-year-old landrace “Mazia” at Areka Agricultural Research Center (Areka, Ethiopia) sequenced in this study. Mazia is one of the least susceptible landraces to Xvm infection [100]. Photos provided by Sadik Muzemil.

Enset is believed to have been domesticated approximately 10,000 years ago in Ethiopia, alongside coffee, various pulses and tef [18]. Often called “the tree against hunger”, highly drought tolerant enset holds historical importance as a year-round source of food for South and South-Western Ethiopia during recurrent famine and drought periods [18]. Enset-based farming systems outperform any other Ethiopian cropping systems in energy yield per area per year [136], and its soil-enhancing and shade-providing qualities make it well-suited for intercropping with root or cash crops like coffee. Cultivation spans diverse agro-ecologies, between 1200 and 3100 m altitudes (up to 3450 m, Muzemil, S. pers. observation) [18], with smallholders growing up to 28 landraces to exploit specific traits including drought tolerance, disease resistance and preferred culinary properties [150].

The extended life cycle of enset presents both advantages and challenges for improvement. Key corm and pseudostem traits are not evident until 2–7 years from seedling establishment, with flowering taking 8–12 years and often resulting in hard, sterile seeds. This is about 2–5 times longer than banana to reach harvest. In mature enset, the pseudostem can reach 0.5–2 m circumference, with up to 17 leaves, each up to 5 m long which nourish an underground corm that can weigh up to 88 kg [152]. These traits likely drove enset’s utilization by prehistoric hunter-gatherer societies [57] and its subsequent domestication. Today enset is a cornerstone of Ethiopia’s economic, social, and cultural fabric [29]. Enset is harvested prior to, or immediately after, flower emergence and before flower maturation while starch reserves are maximal. Because vegetative propagation has been universally adopted, elite landraces are currently identified through time-consuming phenotypic characterization of mature, clonally propagated domesticated landraces over multiple years [152]. Thus, the capacity for beneficial trait selection would markedly benefit from genome-assisted breeding [52].

Enset productivity is threatened by multiple pests and diseases [1, 2, 42, 74], with enset Xanthomonas wilt (EXW) posing a significant threat to food security [154, 155]. EXW is caused by the bacterium *Xanthomonas vasicola* pv. *musacearum* (Xvm) [126], which also causes banana Xanthomonas wilt (BXW) across east and central Africa [13, 103]. Known resistance to Xvm is limited to wild banana (*M. balbisiana*) [102], with only a few genotypes showing some level of tolerance in edible banana germplasm. Importantly, certain enset landraces show reduced susceptibility to Xvm [100, 44], making them popular among enset farmers and thus comprise a considerable proportion of farmyard landraces [153]. These landraces represent a potential genetic resource for enset improvement and BXW resistance breeding in banana. Plant disease resistance is often mediated by nucleotide-binding leucine-rich repeat (NLR) genes [66, 93], yet the NLR repertoire of enset and its relationship to banana NLR families has not yet been assessed. Previous studies have revealed that strong genotype × environment interactions for major agronomic traits [149, 148] have driven regional differentiation of enset landraces [152]. This has resulted in complex vernacular naming systems across ethno-linguistic communities that confound identification of landraces for breeding. Indeed, genome sequencing has revealed significant genetic diversity amongst landraces [151], reflecting local selection and environmental adaptation.

We previously published the first draft genome sequence of *E. ventricosum* [55] and subsequently sequenced 17 Ethiopian genotypes [151], providing initial genomic resources. However, these short-read assemblies had limited contiguity, constraining comparative analyses. The availability of high-quality chromosome-level genome assemblies of closely related species, notably the major diploid progenitors of *Musa* species i.e., *M. acuminata* [9] and *M. balbisiana* [140] or their triploid cultivars [23], provided the opportunity to undertake comparative genomic analyses to identify fundamental differences between these phenotypically similar but agriculturally distinct species. Despite their shared ancestry, the extent of genomic divergence between enset and banana at the gene level, and the biological processes shaped by species-specific gene evolution, remain largely unknown.

Here, we (1) sequenced and annotated the genome and transcriptome of the EXW-resistant/tolerant landrace Mazia (Figure 1C), (2) performed comparative genomic analysis with the main progenitors of diploid banana: *M. acuminata* (MA) [9] and *M. balbisiana* (MB) [140], and (3) characterised the nucleotide-binding leucine-rich repeat (NLR) disease resistance gene repertoire of enset and banana to seek insight into potential tools that may mitigate the major impact of EXW/BXW on these clonally propagated crops. We utilised sequencing data from 20 enset genotypes [151] and 48 banana accessions [9, 140, 23, 115] to assess presence-absence variations and identify genes or gene families unique to enset, providing new insight into their evolutionary roles.

## 2 Materials and Methods

### 2.1 Sample collection

For genomic DNA extraction ∼5g of fresh unexpanded leaf material of 2 year old *E. ventricosum* (EV) landrace Mazia (Figure 1C) was obtained from the national enset *ex-situ* germplasm at Areka Agricultural Research Center (Areka, Ethiopia) and immediately frozen in liquid nitrogen.

For RNA sequencing (RNA-seq), harvested Mazia leaf samples were immediately subjected to stress treatments (cold, heat or wounding) or left untreated before collecting 1-2g of tissue (cut into small pieces) per treatment in RNA*later*^TM^ (Invitrogen^TM^) according to the manufacturer’s instructions. Various other samples, e.g., unexpanded and fully expanded leaves or roots were also cut in small pieces and preserved in RNA*later*^TM^.

### 2.2 Nucleic acid extraction and sequencing

Genomic DNA was extracted using a modified CTAB protocol [62] and used to construct single-tube long fragment read (stLFR) libraries and generate paired-end 100 or 142-base sequence reads on a BGISEQ-500 platform.

RNA extraction was performed according to Logemann et al. [86]. RNA concentration and integrity were assessed using a 2100 bioanalyzer (Agilent) prior to library construction and Illumina sequencing. Briefly, samples from 2.1 were removed from RNA*later*^TM^, quickly blotted dry and immediately frozen in liquid nitrogen. RNA was isolated by guanidine hydrochloride (GuHCl) extraction [86]. Samples ground to a fine powder in a liquid nitrogen were rapidly mixed with GuHCl solution (GuHCl 8 M, MES 20 mM, EDTA 20 mM) and this mixture was extracted with phenol:chloroform (1:1). The resultant aqueous phase was mixed with 0.7 vol. ethanol and 0.05 vol. acetic acid to precipitate the RNA. Following centrifugation, the RNA pellet was washed with 70% ethanol, air-dried and dissolved in sterile water. RNA samples were assessed using a plant RNA Nano assay (Agilent, Bioanalyzer 2100) to determine RNA concentration and integrity and those with a RIN>6.5 were sent to BGI (BGI Health (HK) Company Limited) for library construction and RNA sequencing.

### 2.3 Enset genome assembly and annotation

#### 2.3.1 Assessment of genome size, heterozygosity and ploidy

A *k*-mer (*k* = 21) based analysis was used to estimate the size and heterozygosity of the enset genome. Illumina 2×100 paired-end reads were first trimmed and filtered in TrimGalore (v.0.6.7) (https://github.com/FelixKrueger/TrimGalore) parameter –quality set to 30 and stLFR reads were quality filtered using stLFRdenovo (v1.0.5) (https://github.com/BGI-biotools/stLFRdenovo). The frequencies of 21-mer sequences were calculated with Jellyfish (v2.3.0) [91] from the cleaned reads and used for estimating the genome size and heterozygosity in FindGSE [128] and GenomeScope v2.0 [113], respectively. Smudgeplot (v.4) [113] was used to estimate ploidy of EV genomes using quality filtered stLFR or Illumina paired-end reads.

#### 2.3.2 *De novo* assembly of the enset genome

Adapter sequence filtered stLFR reads were assembled using Supernova (v2.1.1) [141] for stLFR reads (https://github.com/BGI-Qingdao/stlfr2supernova_pipeline#ref). Completeness of the assembled genome was assessed by Benchmarking Universal Single-Copy Orthologs (BUSCO) v.5.2.1 [89] using land plants (embryophyta_odb10) lineage with -long parameter option.

#### 2.3.3 Repeat identification in enset genome assembly

A repeat sequence library was first generated for the enset genome using a combination of homology, *ab initio* and transposable element (TE) structure-based prediction methods. This library was then used to identify repetitive sequences in the enset genome.

For homology-based repeat element prediction, RepeatProteinMask (v.4.1.1) was used with a *p*-value threshold of 1e^-6^ and command-line options -noLowSimple -engine ncbi to search against the RepBase database (RepBaseRepeatMaskerEdition-20181026) at the protein level. Then, RepeatMasker (v3.2) [129] was executed using options -nolow -no_is -norna -engine ncbi -species Viridiplantae on the RepeatProteinMask-masked genome against Dfam [61] v3.2 and the RepBase [70] database for DNA-level transposable element searching.

Subsequently, the genome assembly was further scanned for *ab initio* repetitive element prediction using Miniature Inverted-repeat Transposable Elements (MITE)-Hunter [53], Tandem Repeat Finder (TRF98 v4.09) [10], Piler (v1.0) [35] and RepeatModeler2 (v2.0.1) [38]. TRF was run with 2 5 7 80 10 50 2000 -m -h command-line options, MITE-hunter with -n 5 -S 12345678 –P 1, and PILER as described in https://www.drive5.com/piler/. All predicted *ab initio* repeat elements were merged and processed to exclude sequences containing protein coding genes as determined by BLASTP (e-value threshold of 1e^-10^) against a monocot protein database downloaded from Ensembl Plants (https://plants.ensembl.org/index.html) [16] and Banana Genome Hub (https://banana-genome-hub.southgreen.fr/) [32]. The resulting *ab initio* predicted repetitive elements not encoding proteins were combined with the repeats found by homology. Redundant sequences that did not satisfy the 80-80-80 rule [142] were removed using CD-HIT (v.4.8.1) [83] (parameters -d 0 -c 0.80 -n 5 -G 1 -g 0 -M 0).

For structure-based identification of repetitive sequences, Extensive *de-novo* Transposable Element Annotator (EDTA v1.9.9) [106], a combination of multiple TE predicting tools [105, 121, 127, 147, 157] was used to identify TEs. For EDTA, coding sequences of *Musa* genomes i.e. *M. acuminata* (v4.0) and *M. balbisiana* (v1.1) downloaded from Banana Genome Hub (https://banana-genome-hub.southgreen.fr) and TEs identified by RepeatProteinMask or RepeatMasker were used as input for –cds and –rmout parameters of EDTA, respectively.

TEs predicted from EDTA were finally merged with the non-redundant repeat library obtained from homology and *ab initio*-based predictions. The combined repeat library was supplied to RepeatMasker and used to identify and mask repetitive regions of the enset genome prior to gene prediction. TE families identified by RepeatMasker were further annotated using TEsorter (v1.4) [157].

#### 2.3.4 Gene prediction in enset genome assembly

A combination of *ab initio* gene prediction, homology-based gene prediction and evidence-based gene prediction approaches from BRAKER2 (v2.1.6) [20] and MAKER (v3.01.03) [24] pipelines were applied to annotate gene structure from repeat-masked enset genomes.

BRAKER2 was used in combination with GeneMark-ETP [87, 63, 46, 22] and AUGUSTUS (v3.4.0) [59] to predict *ab initio* gene models supported by protein homology and/or RNA-seq evidence. The raw RNA-seq data from root and young enset leaves of landrace Mazia were aligned against repeat masked genome with hisat2 (v2.1.0) [75] –max-intronlen 160000 –no-mixed –very-sensitive –no-discordant parameter settings, and the alignment processed with SAMtools to generate indexed RNA-seq alignment for input to BRAKER2. Amino acid sequences of *Arabidopsis thaliana* and 11 monocot species were obtained from Phytozome (v13) database and Banana Genome Hub and used as homology-based evidence: *Oryza sativa, Oryza indica, Sorghum bicolor, Dioscorea rotundata, Brachypodium distachyon, Zea mays, Ananas comosus*, *Musa acuminata, Musa balbisiana, Musa schizocarpa,* and *Ensete glaucum*.

Three rounds of MAKER predictions were performed to generate the final enset gene structures. The first two rounds were used to train SNAP (version 2006-07-28) [78] *ab initio* prediction and the third round for combining all the *ab initio*-, homology- and evidence-based gene models. RNA-seq data from roots and leaves of enset landrace Mazia were assembled using Trinity (v2.1.1) [47] with default parameter settings and further analysed to build splice-aware and unique transcript assemblies using PASA pipeline (v2.4.1) (https://github.com/PASApipeline/PASApipeline). The assembled Mazia transcripts, together with proteins from *A. thaliana* and the 11 monocots, were imported into MAKER as evidence and homology-based hits used to generate the initial gene models using MAKER settings est2genome=1 and protein2genome=1 with no *ab initio* gene predictors. SNAP was then trained with gene models from this first-round prediction and the resulting gene models were added as *ab initio* input to MAKER for second round prediction. These were then used to retrain and generate the final SNAP *ab initio* gene models for MAKER. Evidence for protein coding was determined by integrating SNAP, GENEMARK and AUGUSTUS gene models, in combination with protein homology and evidence from assembled RNA-seq transcripts in the MAKER pipeline. High-confidence gene models having an Annotation Edit Distance (AED) of less than or equal to 0.25 were retained as a final gene model set of MAKER annotation. AED [36], a gene annotation quality measure in MAKER assesses the strength of agreement between a gene annotation and the supporting evidence supplied for gene prediction.

To select the final set of gene models, the predicted protein coding gene models (AED < 0.25) were assessed for inserted TEs that might not have been detected by similarity or TE structural feature detecting tools in the repeat identification and masking steps. Differential TE insertions are prevalent in crop genomes [139] highlighting the need for manual, semi and/or automated curation of structurally annotated genes. Hence, to identify TEs containing predicted protein coding genes (AED < 0.25) of enset, we used TEsorter (v1.4) [157] with parameters -hmm rexdb-plant –pass2-rule 80-80-80 and excluded any gene models harbouring TEs and ran BUSCO to assess annotation completeness on the remaining predicted genes.

#### 2.3.5 Functional annotation of predicted protein-coding genes

Functional annotation was performed using sequence-similarity searches against the NCBI non-redundant (NR) and the Clusters of Orthologous Groups (COGs) databases [131], SwissProt, and TrEMBL [15] using DIAMOND’s [21] BLASTP (e-value < 1e^-5^, –ultra-sensitive) and Evolutionary Genealogy of Genes: Non-supervised Orthologous Groups (eggNOG)-mapper (v2.1.7) [25] programs with default parameter settings. In addition, local InterProScan (v5.52-86.0) [14] was performed to scan for motifs and domains and ascribe Gene Ontology (GO) terms for annotated genes, and BUSCO was used to assess completeness of the predicted proteome. The GO terms were filtered and synchronised with the updated GO terms and in the core GO to retain their description and exclude obsolete terms (http://geneontology.org/docs/download-ontology/).

#### 2.3.6 Annotation of non-coding RNA (ncRNA) genes

Enset genomes were scanned for tRNAs using tRNAscan-SE (v 2.0.9) [28] with eukaryotic parameters. Ribosomal RNA (rRNA) fragments were predicted by BLASTN (e-value < 1e^-5^) homology search of *Arabidopsis* (5S, 5.8S and 18S rRNA) and rice (28S rRNA) template rRNA sequences against the assembled genome. MicroRNA (miRNA) and small nuclear RNA (snRNA) sequences were identified using INFERNAL (v1.1.4) [104] cmscan with default parameter settings against the Rfam [71] database as a reference. The Rfam library of covariance model from (ftp://ftp.ebi.ac.uk/pub/databases/Rfam/14.8/Rfam.cm.gz) and clanin file from (ftp://ftp.ebi.ac.uk/pub/databases/Rfam/14.8/Rfam.clanin) were supplied to cmscan for the detection of miRNA and snRNA sequences.

### 2.4 Comparative analysis of enset and banana genome sequences

#### 2.4.1 Genome assemblies data

In addition to sequence data generated during this study, we obtained data from our previous genome assembly (GenBank: GCA_000818735.3) from the GenBank [118] database via the National Center for Biotechnology Information (NCBI) [117] web portal. We also downloaded *M. acuminata* (v4.0) and *M. balbisiana* (v1.1) reference genome assemblies from the Banana Genome Hub (https://banana-genome-hub.southgreen.fr) [32].

#### 2.4.2 Sequence reads data

Genomic sequence reads for 19 EV and 48 banana (*Musa* spp.) genotypes were obtained from NCBI Sequence Read Archive (SRA) [77] (Supplementary Tables 2 and 3) database using esearch (v16.2) and converted into FASTQ format in fastq-dump (v.3.03 options: –split-files –gzip –skip-technical). Long reads of *Musa balbisiana* were obtained from CNGBdb with ID number CNR0255329 (https://db.cngb.org/search/run/CNR0255329/).

For genomic reads mapping, a total of 370 Gb of NGS reads from 20 enset genotypes were utilised, representing a total of 650 X coverage of the enset haploid genome. We used 942.9 Gb (1800 X) of sequence reads from 36 Musa AA-genome (AA or AAA type) genotypes, and 418.8 Gb (974 X) from 12 Musa BB-genome (BB-types). These sequence reads, including both short and long reads, are listed in Supplementary Tables 2 and 3.

In total, including the data generated in this study, enset NGS reads cover 16 cultivated landraces (including Arkiya, Bedadeti, Buffero, Derea, Lochinge, Mazia, Nechiwe, Nobo, Siute and Yako) and 2 wild genotypes (Erpha 13 and Erpha 20) originally obtained from enset growing areas in South and Southwestern Ethiopia [151]. However, two genotypes (ITC1387 and JungleSeeds [55]) had unknown origins (Supplementary Table 2).

The banana genotypes were double haploid (DH)-Pahang (AA), Pisang Kra (AA), Pisang Mas (AA), Green-Red (AA), Calypso (AA), subsp. Zebrina (AA), Cavendish (AAA), Gros Michel (AAA), DH-Pisang Klutuk Wulung (BB), Cultivar DYN018 (BB), Cultivar DYN049 (BB), Cultivar DYN019 (BB) and Cultivar DYN302 (BB) (Supplementary Table 3).

#### 2.4.3 Quality control for sequence reads data

Downloaded Illumina paired reads were assessed for quality using FastQC (v0.11.9) [4]. Reads were trimmed on quality score and filtered for adapter sequences using TrimGalore (v.0.6.7) (https://github.com/FelixKrueger/TrimGalore, parameter: –quality 30). The stLFR paired-end reads of landrace Mazia generated in this study were quality filtered using stLFRdenovo (v.1.0.5) (https://github.com/BGI-biotools/stLFRdenovo).

Pacific Bioscience high-fidelity (PacBio HiFi) long sequencing reads [84] were also cleaned from adapter sequences using cutadapt v.4.3 [92] with the following pa-rameters: -b “ATCTCTCTCAACAACAACAACGGAGGAGGAGGAAAAGAGAGAGAT;min_overlap=35” -b “ATCTCTCTCTTTTCCTCCTCCTCCGTTGTTGTTGTTGAGAGAGAT;min_overlap=35” –discard-trimmed –revcomp -e 0.1 –report=minimal –discard. PacBio contiguous long reads (CLRs) were error-corrected using Canu v2.3 and Porechop v0.2.4 (https://github.com/rrwick/Porechop, parameter: –adapter_threshold 95) was used to remove adapter sequences from Oxford Nanopore Technology (ONT) reads of banana genotypes (*Musa* spp.).

#### 2.4.4 Aligning sequence reads versus reference genome assembly

Filtered and trimmed *E. ventricosum* (EV) NGS reads were mapped to *M. acuminata* (MA, AA-genome) and *M. balbisiana* (MB, BB-genome) genome assemblies using bwa-mem2 [138]. Non-aligned reads were discarded using SAMtools (v1.22) (parameter: -F 4) and PCR duplicates were filtered out using the MarkDuplicates package of PICARD tools (v2.27.1) [19]. Similarly, quality filtered NGS reads of *Musa* AA-genome (AA or AAA type) genotypes and *Musa* BB-genome (BB-type) genotypes were mapped against the two EV genome assemblies, landraces Mazia and Bedadeti. In addition, NGS reads of EV genotypes were also mapped back to EV genome assemblies, AA or AAA type banana genotype reads were mapped to the MA genome, and BB-type reads were mapped to the MB genome.

For long read mapping of banana genotypes against EV assemblies, minimap2 (v 2.24-r1122) [82] with options -ax map-hifi, -ax map-pacbio, and -ax map-ont was used to map error corrected or adapter sequence trimmed PacBio HiFi reads, PacBio CLR reads and ONT reads, respectively. These long reads of banana genotypes were also mapped back to genome assemblies of their respective genome groups as described for NGS reads.

#### 2.4.5 Coding sequence (CDS) alignment

Prior to CDS alignment between genomes of EV, and *Musa* spp. (MA and MB), repetitive sequences were first hard-masked with BEDtools (v2.30.0) [109] maskfasta function using genomic coordinates of non-genic regions. In addition, CDS that showed greater than 25% interspecies genomic reads mapping coverage were discarded (see Section 2.4.4 for details).

Next, we aligned CDS of enset genomes against CDS of MA and MB genomes using NUCmer [90] and LASTZ [54]. NUCmer (MUMmer4 v4.0.0rc1) [90] alignment was generated first and this was parsed into coordinate representation of the aligned genomic position using show-coords (parameters: -c -d -l -r -T) in MUMmer4 [90]. LASTZ [54] was run on CDS that showed < 25% NUCmer alignment coverage setting parameters –strand=both –gapped –format=PAF options. CDS with less than 25% LASTZ alignment coverage were retained for further analysis. Scripts used for processing the alignments are available at https://github.com/sadikmz/EV_MAB_compGeno.

#### 2.4.6 Overview of approach to finding species-specific genes in enset and banana

To assess the similarity between enset (*E. ventricosum*) versus the major diploid progenitors of cultivated banana (*Musa* spp.), we performed coding sequence (CDS) alignments both at the level of assembly versus assembly and also at the level of sequence reads versus assembly. We utilised genome sequencing reads and genome assemblies of *E. ventricosum* (EV), *M. acuminata* (MA), *M. balbisiana* (MB) and triploid banana (AAA).

Two approaches were used to determine conservation of genes between enset and banana, and thus presence/absence variations (PAVs). The first involved aligning CDS from annotated genome assemblies of one species against CDS from another. The second used alignment of genomic sequence reads from one species against another. In both approaches we examined the fraction of the gene covered by the alignment (breadth of sequence alignment) and where CDS showed less than 25% coverage, we considered that CDS (and its corresponding gene) to be absent. This enabled identification of species-specific genes that might have evolved *de novo,* been lost in one of the species or undergone divergence from the CDS of their last common ancestor.

#### 2.4.7 Inferring presence/absence of genes from alignment coverages

Genome-wide gene coverage of mapped genomic reads was extracted using BEDtools (v2.30.0) [109] coverage parameter. The obtained coverage values, along with merged LASTZ and NUCmer alignment coverage across coding sequences, were used to identify candidate presence-absence sets of genes among the enset and banana genomes. Breadth of CDS coverage from alignment and reads mapping, and read depth extracted from primary alignments was visualised in karyoploteR [43] R package and Jbrowse [31], respectively. Read depth of primary alignment was extracted using deepTools [111] bamCoverage. Qualimap (v.2.2.2-dev) [41] was used to summarise mapping statistics.

#### 2.4.8 Functional annotation of differentially present/absent genes

Identified candidate EV, MA and MB specific genes were assessed for Gene Ontology (GO) [3] terms associated with “Biological” terms. Integrated Genome Viewer (IGV) [133] v2.11.9, tidyverse [143] and ggcoverage [123] R packages were used for data visualisation. Genes of presence-absence variation in enset and *Musa* species that lacked GO-terms annotation were further searched against NCBI NR and SwissProt curated proteins databases and the proteomes of *Arabidopsis thaliana* (TAIR10) and *Oryza sativa* (v7.0) obtained from Phytozome (v13) using BLASTP (e-value < 1e^-5^). In addition, InterProScan (v5.52-86.0) [14] and Proteinfer [116] scan was performed to detect gene motifs and conserved domains in the amino acid sequences.

#### 2.4.9 Clustering of proteins into orthogroups

EV, MA and MB specific protein-coding genes that lack detectable homology inferred from sequence similarity with proteins of closely or distantly related species were used to cluster them into orthogroups. Clustering proteins into orthogroups was achieved using OrthoFinder (v.2.5.5) with parameters -M msa -A muscle -T raxml. The resulting output was used for protein structure prediction.

#### 2.4.10 Predicting protein structure

The 3D structures of individual proteins were predicted using AlphaFold2 [68] in monomer model. High-confidence predicted structures showing predicted local-distance difference test (pLDDT) > 70% were further assessed with Foldseek [7, 76, 137] (https://search.foldseek.com/search) to infer structural similarity with other proteins. The most significant hits (with a *p* value < 0.01) against computationally or experimentally validated structures from protein database were retained. These were visualised with ChimeraX v1.8.

### 2.5 NLR encoding genes and phylogenetic analysis

#### 2.5.1 Identification of NLR encoding genes in genome assemblies

Genomic locations harbouring nucleotide binding domain leucine rich repeat (NLR) genes in *E. ventricosum* (EV) landraces Mazia and Bedadeti, *Musa acuminata* (MA) and *Musa balbisiana* (MB) genomes were assessed using NLR-Annotator [125] using default parameter values. Predicted NLR loci domains were annotated using the annotateClasses.py Python script [8]. Subsequently, nucleotide sequences of predicted NLR loci in EV were subjected to a reciprocal BLASTN search (e-value < 1e^-5^) with enset against *Musa* genomes, and vice versa, to identify their homology.

In addition to searching for NLRs in the genomic DNA sequences, we also searched for NLRs among the amino acid sequences of predicted protein coding genes in the EV, MA and MB genomes. Briefly, a local database of NLRs from a diverse range of plants was created using predicted and experimentally validated NLRs in ANNA [85] (https://biobigdata.nju.edu.cn/ANNA/) and RefPlantNLR [80], respectively. The protein sequences of enset and *Musa* species genomes were then searched against the local NLR database using DIAMOND BLASTP (e-value < 1e^-5^ –ultra-sensitive) and hits were then aligned in Clustal Omega (1.2.1) [122] with default parameter values to build Hidden Markov Models (HMM) using hmmbuild of HMMER [58] (v.3.3.2). The HMM from the local NLR database was subsequently added to the list of NLR-associated Pfam domains [14]: NB-ARC (PF00931), TIR (PF01582, PF13676), RPW8-type (PF05659), Late blight resistance protein R1 (PF12061), Rx-type (PF18052) and LRRs (PF00560, PF13516, PF18805, PF13855, PF07725, PF12799, PF01463, PF01462, PF18831, PF18837, PF08263, PF07723, and PF13306). The combined HMMs for NLRs were then used to scan against proteins of EV, MA, and MB genomes with hmmsearch (-E 0.0001 and –domE 0.001) in HMMER (v.3.3.2). The identified NLR genes were rescanned for the NLRs highly conserved NB-ARC (PF00931) Pfam domain with hmmsearch using the same parameters to confirm their presence in each sequence. Domains detected in NLR genes were annotated with NCBI Conserved Domains Database (CDD) and InterProScan run with default parameters to identify coiled coil (CC), NB-ARC (N) and RPW8 type domains. The LRRs 1, 9 and 11 MEME (Multiple EM for Motif Elicitation) motifs [69] were used to detect LRR domains of NLR genes using Motif Alignment and Search Tool (MAST) with default parameters.

The identified NLR genes in each genome were also assessed for their overlap with NLR loci predicted by NLR-Annotator from the same genome. NLR genes were mapped back to their individual genomes using TBLASTN (e-value < 1e^-5^) and overlapping NLR loci from the two sources were compared using the intersect function in BEDTools (v2.30.0) [109] and a custom R script. NLR loci without a corresponding predicted protein were subjected to BLASTX (e-value < 1e^-5^) against the NCBI NR protein database to detect homology to NLRs in other species, thereby increasing the confidence they encoded genuine NLRs.

#### 2.5.2 Phylogenetic analysis of NLRs

The core NB-ARC domains NLRs encoded in genomes of EV landraces Mazia and Bedadeti, MA and MB were aligned using Clustal Omega [122] and all positions with gaps > 50% in aligned sequences were removed using trimAI [26]. The evolution of NB-ARC domains of NLRs was inferred with RAxML (v8.2.12) [124] using the Maximum Likelihood method based on the JTT model and a bootstrap value of 1000 replicates. Non-plant NB-ARC containing domains from RefPlantNLR [80] were included as outgroups to root the phylogenetic tree. The resulting tree was visualised using the Interactive Tree of Life (iTOL) [81].

## 3 Results

### 3.1 The enset Mazia genome is triploid

We sequenced the genome of EXW-tolerant landrace Mazia using stLFR at 330X coverage. We estimated the Mazia genome size to be 572.23 Mb and heterozygosity at 1.41%, using the frequency distribution of 21-mers from these unassembled quality-filtered stLFR reads (Figure 2A and Supplementary Figure 1). K-mer-pairs of k=21 differing by one SNP showed a strong signal of a triploid Smudgeplot pattern (Figure 2B) near to 0.3 normalized coverage indicating the Mazia genome is triploid.

**Figure 2:**
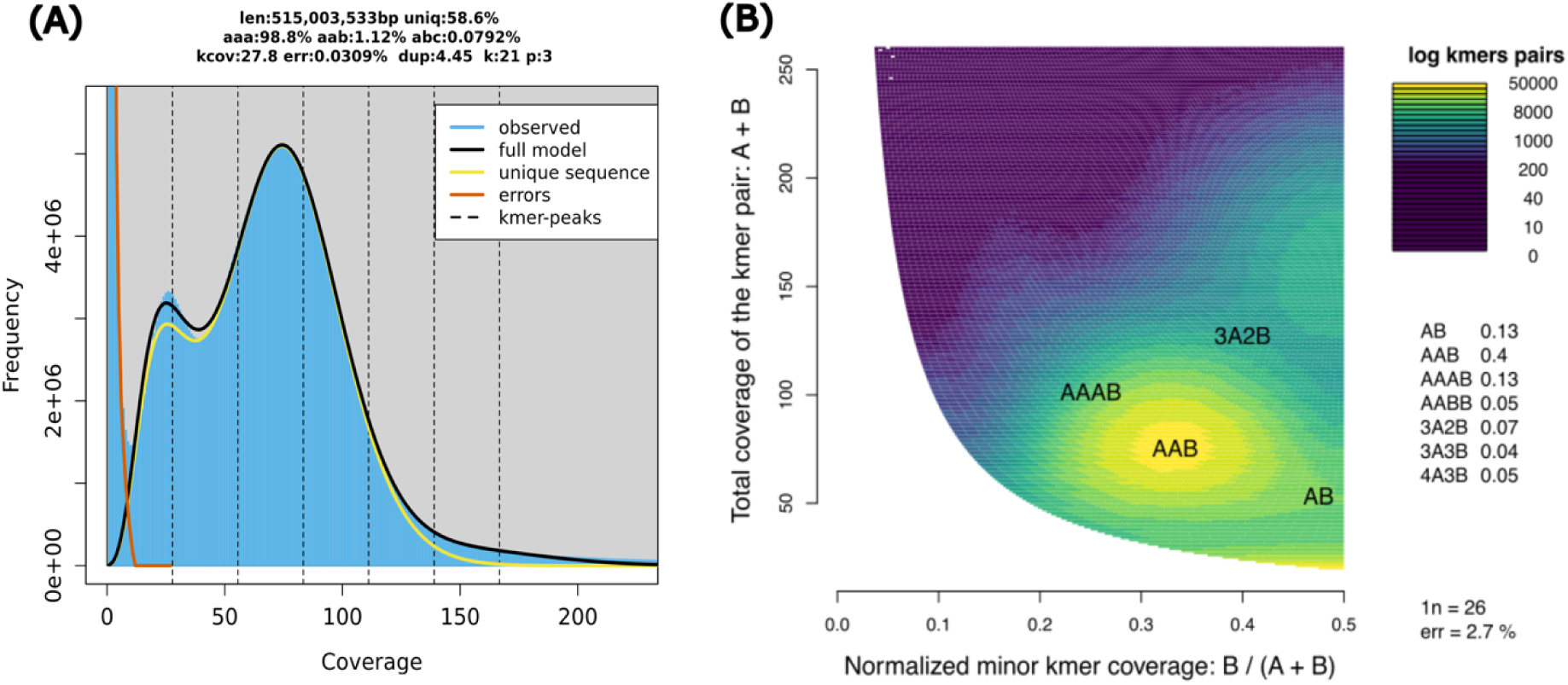
Kmer-based inferred genome profile of *E. ventricosum* (EV) landrace Mazia. (A) GenomeScope v2.0 [113] inferred characteristics of genome structure, including genome size, abundance of repetitive elements, and rate of heterozygosity of EV Mazia genome. (B) Smudgeplot (v.4) [113] (k=21) inferred ploidy level of EV landrace Mazia suggests it is triploid.

### 3.2 Genome assembly and annotation

The haploid genome assembly length for landrace Mazia was 540.14 Mb, which is approximately 94% of the k-mer based estimated genome size. The assembly comprises 43,459 scaffolds with an N_50_ of 290 kb. The lengths of the scaffolds ranged between 0.5 kb and 2.67 Mb, with an average size of 12.43 kb (Table 1). This represents a substantial improvement over our previous haploid assembly of landrace Bedadeti (GenBank: GCA_000818735.3), which had a total length of 451.28 Mb and an N_50_ of 21 kb.

**Table 1:** Assembly and annotation metrics for enset (*E. ventricosum*) genome assemblies of landraces Mazia and Bedadeti.

| Description | Mazia | Bedadeti* |
| --- | --- | --- |
| **Assembly** |  |  |
| Number of scaffolds | 43,459 | 45,745 |
| Total length of the assembly (Mb) | 540.14 | 451.28 |
| Scaffolds N<sub>50</sub> (Kbp) | 290 | 21 |
| Scaffolds L<sub>50</sub> | 436 | 6010 |
| Length of largest scaffold (Mb) | 2.67 | 0.21 |
| Average length of scaffolds (kb) | 12.43 | 9.87 |
| BUSCO (genome mode S:D in %) | 82.6 : 5.9 | 80.7 : 3.5 |
| Repetitive sequences content (%) | 64.64 | 51.70 |
| **Structural and functional annotation** |  |  |
| Number of predicted protein coding genes | 38,940 | 38,623 |
| Predicted gene length (mean:median bp) | 3591 : 1981 | 3372 : 1890 |
| CDS length (mean:median bp) | 230 : 144 | 229 : 142 |
| Number of exons per spliced gene (mean:median) | 5.8 : 4 | 5.6 : 4 |
| BUSCO (proteins mode S:D** in %) | 68.0 : 5.5 | 68.5 : 3.8 |

GenBank: GCA\_000818735.3; protein coding genes were annotated using RNA-seq and the pipeline employed for landrace Mazia in this study.
S = Complete and single-copy Benchmarking Universal Single-Copy Orthologs (BUSCO) [89]; D = Complete and duplicated BUSCO.

The Mazia genome assembly had a repetitive-element content of 64.64%, compared with 51.70% for the Bedadeti genome assembly (Supplementary Table 1). We identified 38,940 and 38,623 high-confidence gene models for Mazia and Bedadeti respectively, after excluding genes containing TE insertions (Supplementary Table 4). This represents gene densities of 13.87 kb and 11.68 kb, respectively, per gene. These values are comparable to the gene densities of banana genomes, *M. acuminata* (14.12 kb) and *M. balbisiana* (14.80 kb). We assigned putative function to 83% of predicted protein sequences (Supplementary Figure 2) in both Mazia and Bedadeti using NCBI’s non-redundant protein sequence database. For Mazia and Bedadeti we annotated 141 and 137 ribosomal RNAs (rRNAs), 1634 and 1461 transfer RNAs (tRNAs), 212 and 251 micro-RNAs (miRNAs), and 209 and 197 small nuclear RNAs (snRNAs), respectively (Supplementary Tables 11 and 12).

### 3.3 Most, but not all, banana genes are conserved in enset

We next investigated the conservation of banana genes in enset by aligning enset genome sequencing reads against the reference genome sequences of the major diploid progenitors of cultivated banana: *Musa acuminata* (Figure 3A) and *M. balbisiana* (Figure 3B). To complement this approach, we also aligned predicted enset protein-coding DNA sequences (CDS) against the banana reference genomes (Supplementary Figures 3 and 4). Overall, there is a broad conservation of enset and banana genes across both banana reference genomes with less conservation around the centromeres (Figure 3). Of the enset sequence reads, 96.7% and 97.7% were aligned against the MA and MB reference genomes (Supplementary Table 5), covering respectively, 64.7% and 69.6% of the MA and MB genomes’ lengths.

**Figure 3:**
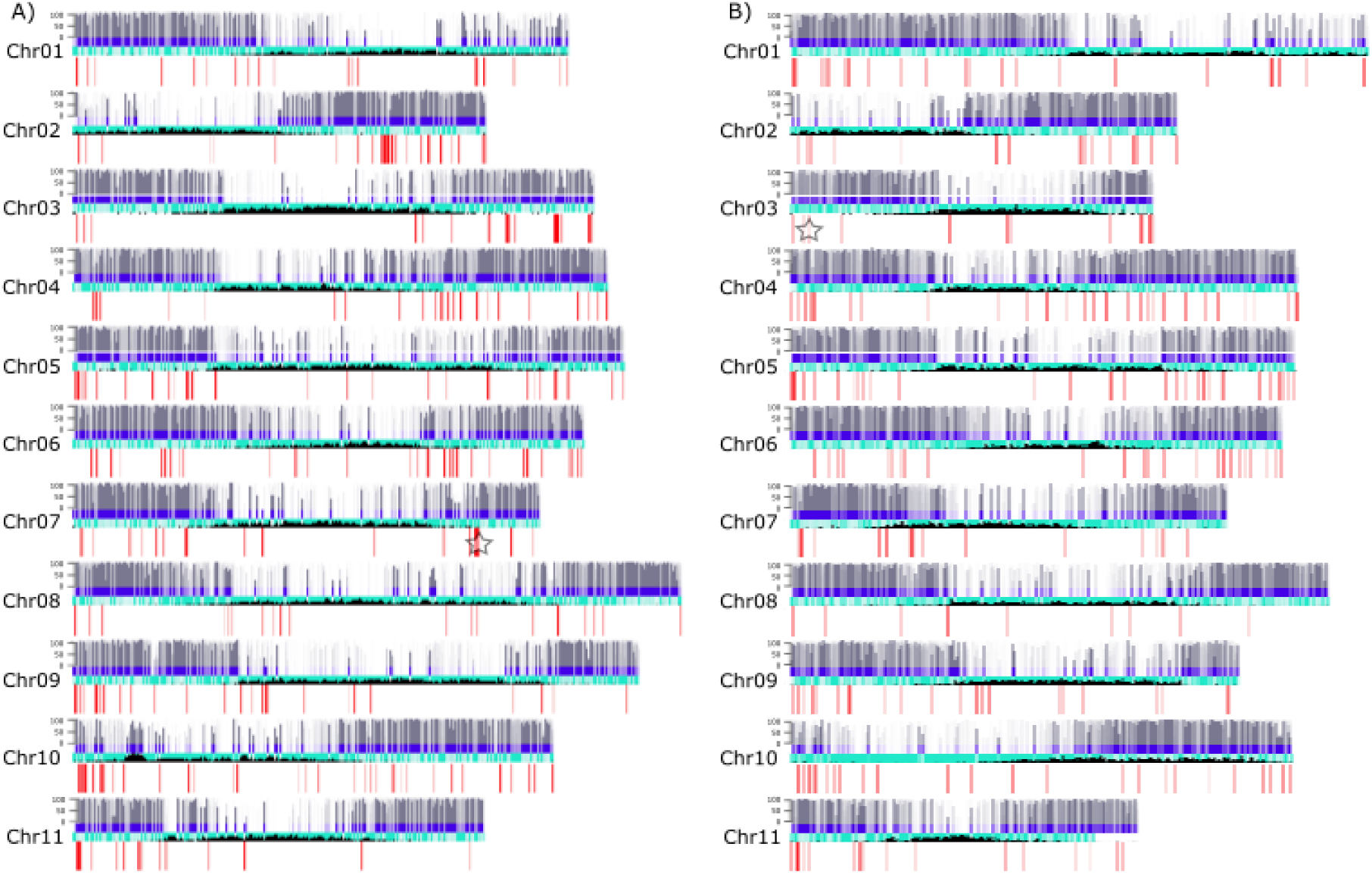
Sequence conservation between banana and enset. Genomic sequence reads of 20 *E. ventricosum* (EV) genotypes were mapped against *M. acuminata* (MA) and *M. balbisiana* (MB) genomes. EV coding sequences (CDS) were aligned against the MA and MB genomes. Breadth of coverage of CDS alignments and reads mapping across MA and MB genes were plotted for MA (A) and for MB (B). Reads mapped with bwa-mem2 (v.2.2.1) default parameter, LASTZ [54] and NUCmer (MUMmer4) [90] were used to produce CDS alignment. Gray coloured bar lines in the upper lane indicate breadth of coverage CDS alignment or NGS reads mapping across each gene (represented by a purple short bars). The grey bar height is proportional to the alignment or mapping fraction coverage per gene locus. Red bar lines underneath indicate gene loci that are absent from enset (<25% breadth of coverage by reads and genomic alignment). Light blue coloured lines below the blue coloured bars represent non-coding regions and black bars indicate the density of centromeric repeats in MA and MB genomes [9]. The stars on MA Chr07 and MB Chr03 indicate the genomic locations specific disease resistance encoding genes (See Figure 4 for details).

Most banana genes are conserved in enset. However, 875 (1%) MA and 615 (1%) MB genes were absent from enset based on the alignment of genome sequencing reads and assembled CDS. Of these, only 314 (MA 35.90%) and 413 (MB 67.15%) were annotated with GO-terms. GO-terms associated with Biological Process for these banana-specific genes included disease response (24 in MA and 8 in MB), regulation of DNA-templated transcription (19 in MA and 12 in MB) and protein phosphorylation (18 in MA and 10 in MB) (Figure 4, Figure 6B and Supplementary Table 9). Additionally, genes involved in abscisic acid-activated signalling pathways (4 in MA and 4 in MB) and the tricarboxylic acid cycle (3 in MA and 1 in MB) were also identified.

**Figure 4:**
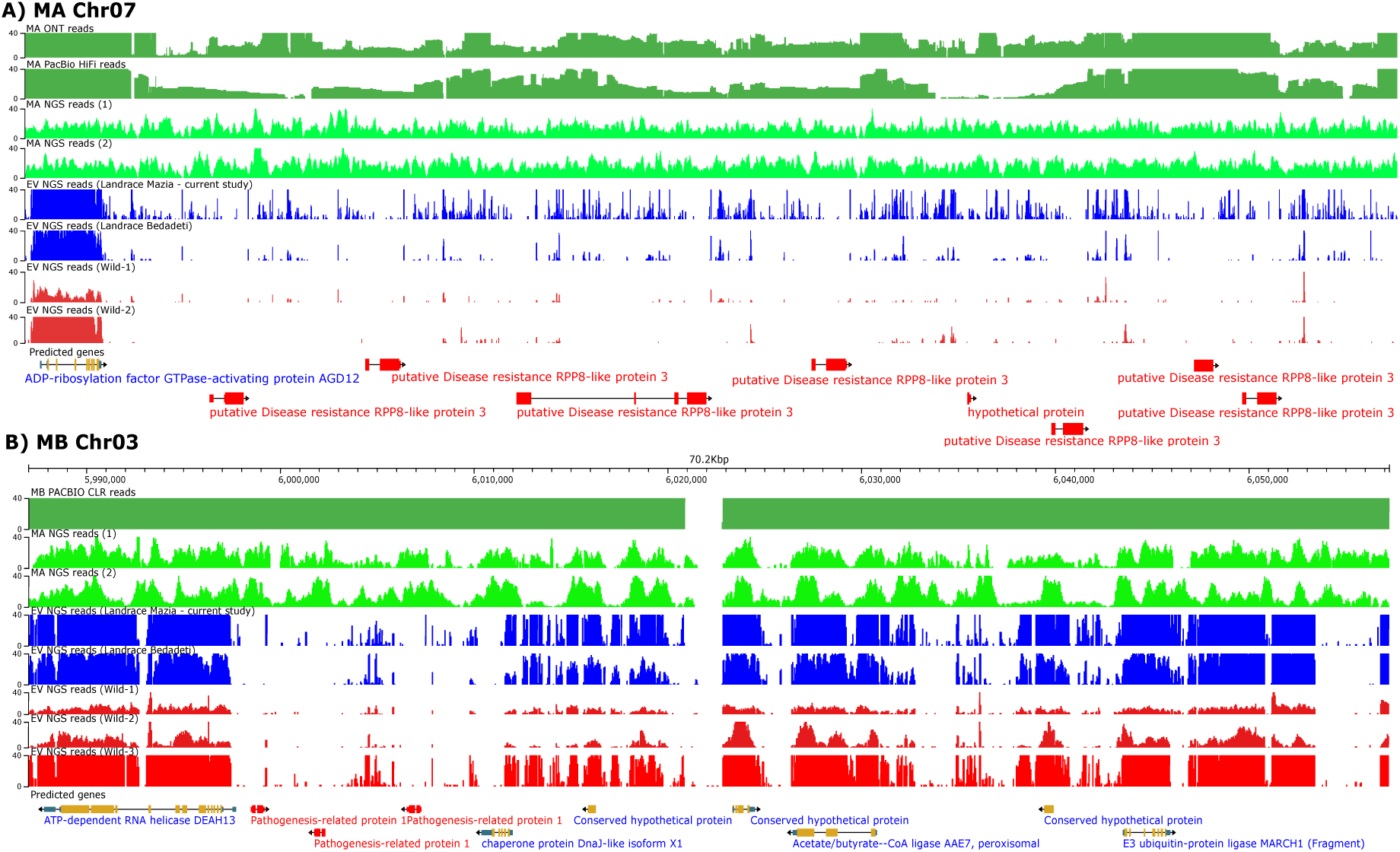
Examples of genomic regions from *M. acuminata* (MA) and *M. balbisiana* (MB) chromosomes protein coding genes absent in landrace Mazia and Bedadeti. (A) A 157 kb region in chromosome 7 of MA, and (B) a 70.2 kb region in chromosome 3 of MB containing MA and MB species specific CDS encoding disease resistance genes that are absent in enset. The first top two lanes of dark green bars in (A) indicate read depths of MA ONT [9] and PacBio HiFi [84] long reads, and the two light green bars represent NGS reads of MA genotypes [SRA: ERR3606950 and ERR3606951]. For (B), the top dark green bar and the subsequent two lanes with light green bars represent NGS read depth of MB genotypes [SRA: ERR10695603]. The blue lanes for both (A) and (B) represent mapping depth of EV NGS reads from domesticated landrace Bedadeti [151] and Mazia (generated in this study), while red lanes indicate NGS reads from wild EV landraces reported by [151] and [55], respectively. Genes are represented in the bottom lane by horizontal lines with yellow boxes and black arrows. MA and MB specific species genes are indicated by horizontal lines containing red bars in the bottom lane of (A) and (B), respectively.

### 3.4 Enset genes that are not conserved in banana

To ascertain which predicted enset genes are conserved in banana, we aligned banana genomic reads, and banana predicted CDS, against the Mazia and Bedadeti enset genome assemblies. The banana sequence data (Supplementary Table 3) included both short reads, and PacBio/ONT long reads for MA [84] and MB [140] (Figure 5A and 4B). About 94.50% of the AA-genome reads and 95.30% of the BB-genome banana genomic reads aligned to Mazia. The banana AA-genome reads covered 65.01% (Mazia) and 68.99% (Bedadeti) of the enset genome assemblies, whereas the BB-genome reads were slightly lower (59.93% in Mazia and 65.71% in Bedadeti) (Supplementary Tables 6 and 7). On average, 93% of the CDS in enset genomes were covered (>25% of their breadth) by aligned banana reads and/or banana CDS. However, 2747 Mazia and 1104 Bedadeti gene predictions did not align against the banana sequences, identifying these as enset-specific genes.

**Figure 5:**
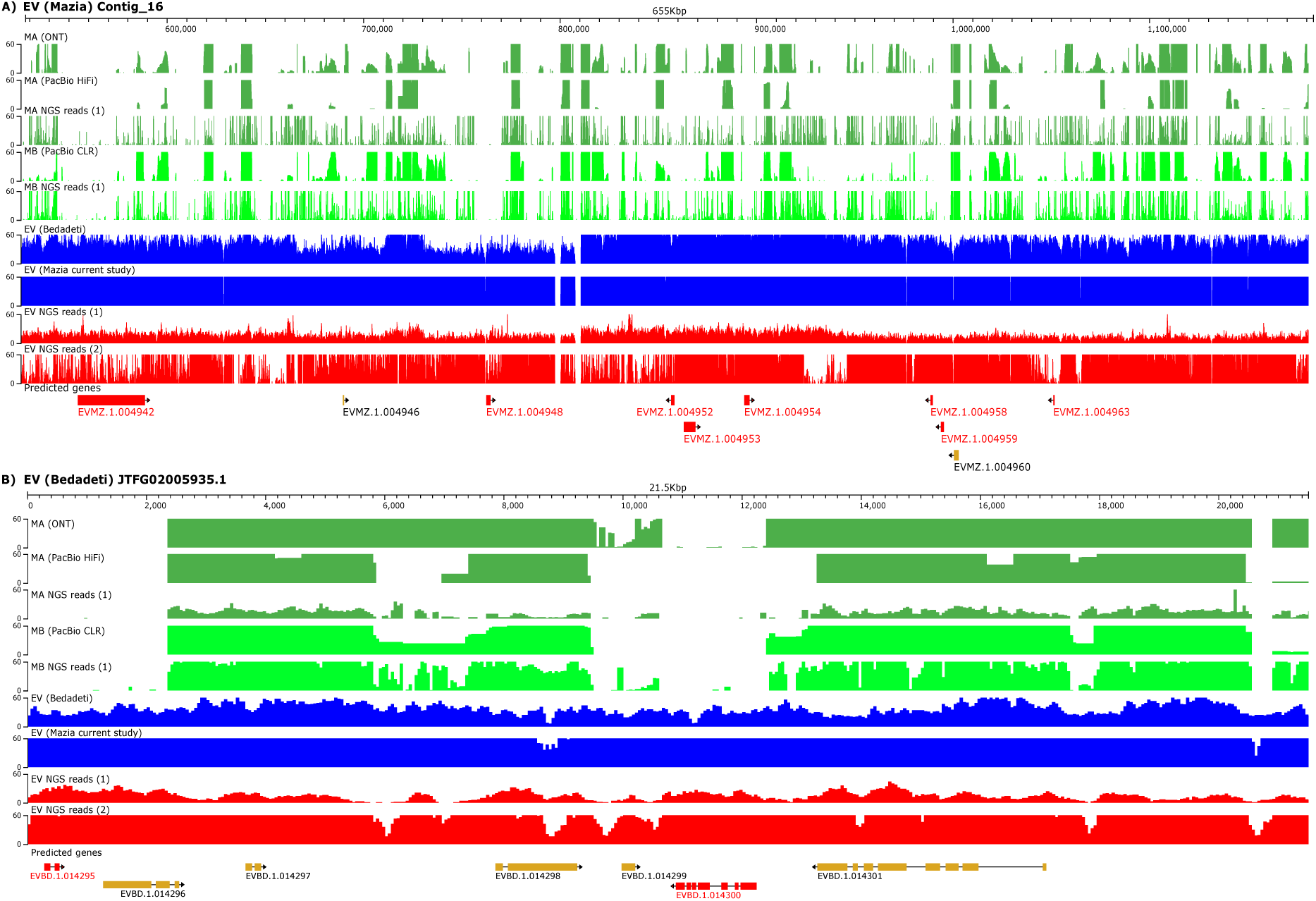
Enset (*Ensete ventricosum*) genomic regions with enset-specific genes. (A) A 655 kb region of EV landrace Mazia genome encodes predicted protein-coding genes that lack coverage by read mapping from *M. acuminata* (MA) and *M. balbisiana* (MB) genomes. (B) A 21.5 kb region of the genome of EV landrace Bedadeti contains enset-specific protein-coding genes. In (A) and (B), from top to bottom, the dark green coloured bars indicate mapping depth of MA ONT [9], PacBio HiFi [84], and NGS reads [SRA: ERR3606950]. The light green coloured bars were for MB PacBio [140] or NGS reads [SRA: ERR10695603], as annotated. The blue lanes represent mapping depth of EV NGS reads from domesticated landraces Bedadeti and Mazia (generated in this study), while red lanes from top to bottom indicate NGS reads from wild EV landraces reported by [151] and [55]. Predicted protein coding genes are indicated in the bottom lane as yellow and red boxes, with red boxes representing an EV specific set of genes that lack either reads mapping or CDS alignment from AA-genome and BB-genome banana sequences.

Our analysis identified enset-specific genes whose GO term annotations were predicted to be involved in a range of biological processes, including DNA integration (2 genes), carbohydrate metabolic process (4 genes) and DNA-templated transcriptional regulation (7 genes) (Figure 6A, and Supplementary Table 8).

**Figure 6:**
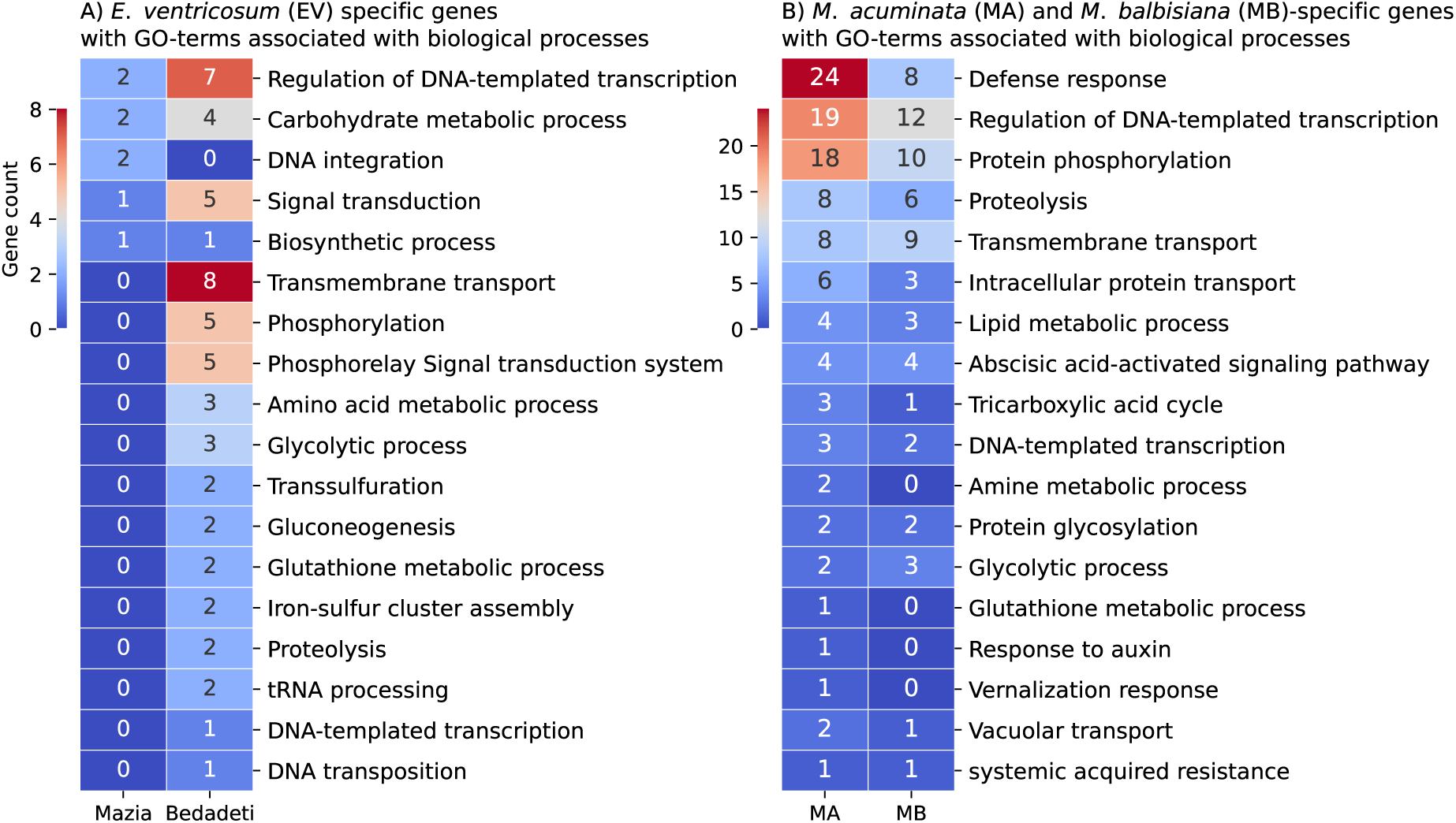
Annotated Biological Processes Gene Ontology (GO) terms for unique genes in Enset or MA and MB genomes. GO terms associated with biological processes in species-specific protein coding genes of EV landraces Mazia and Bedadeti (A), or MA and MB (B) specific protein coding genes. The abundance of identified genes with associated GO terms is indicated by the colour key, with dark blue representing zero and dark red the largest value depicted in A and B. Presence or absence of genes in the three species were identified based on interspecies genome alignment or read mapping across coding sequences. Genes with breadth of coverage <25% of the CDS’s length were considered species-specific (see Supplementary Tables 7 and 8 for detailed lists).

However, approximately 1.29% (502 out of 2747 in Mazia) and 0.97% (375 out of 1104 in Bedadeti) of enset genes lacked sequence homology with genomic reads or CDS in banana AA-and BB-genomes and had no functional annotation. These unique genes most likely evolved for roles in enset-specific processes. Notably, the remaining identified enset-specific genes had predicted functions. These include four proteins associated with TEs, a homolog of disease resistance RPP8 (*Resistance to Peronospora Parasitica 8*) gene, now classified as a “helper” resistance gene, and zinc finger protein transcription factors (Supplementary Tables 13 and 14).

### 3.5 Unique genes

About 1.13% of the EV genes (502 in Mazia and 375 in Bedadeti) not found in MA and MB lacked detectable homology (inferred from sequence similarity with proteins of closely related or distant species) with proteins of closely related or distantly related species. In *Musa* 0.61% (225 out of 36979) of MA and 0.40% (131 out of 33021) of MB genes absent in EV lack detectable homology with proteins of other species. We refer to these sets of genes in enset and banana genomes as “orphan genes” [107, 132, 73, 34] or “unique genes”.

To examine evolutionary relationships of these unique protein encoding genes in enset and banana with proteins in distantly related species we used *ab-initio* prediction of protein structure, allowing homology detection between distantly related proteins lacking sequence similarity [72]. Orphan genes in enset and banana genomes were clustered into orthogroups (OrthoFinder) and the structure of species-specific orthogroup proteins were predicted using AlphaFold2 [68] in monomer model. High-confidence structures showing the predicted local-distance difference test (pLDDT) > 70% were further assessed in Foldseek [7, 76, 137] to infer structural similarity with other proteins.

AlphaFold2 predicted high-confidence (pLDDT > 70%) protein structures for only 14 (out of 877 combined Mazia and Bedadeti) EV, 28 MA (out of 225) and 2 MB (out of 131) unique proteins (Supplementary Table 15). Two of the 14 unique EV proteins showed a very high-confidence structure (pLDDT > 90%) and most showed significant (e-value < 1e^-3^) structural similarity with experimentally determined structures from proteins of unknown functions (Figure 7A and 7B) or predicted structures from enset proteins.

**Figure 7:**
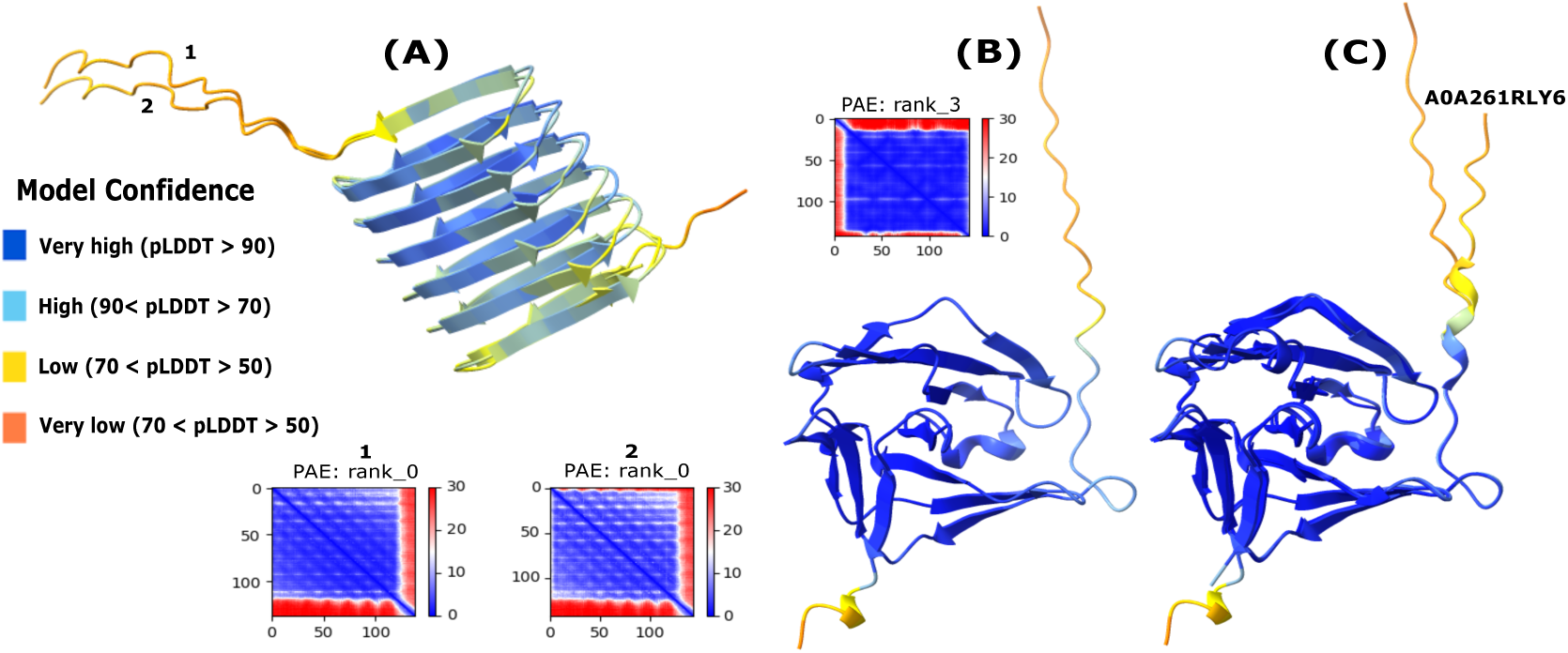
Predicted 3D structures of selected enset unique proteins (*Ensete ventricosum*) and their similarity with distantly related proteins. (A) Aligned protein structures of a pair of unique enset genes from Mazia (#1) and Bedadeti (#2) landraces. (B) A very high confidence (pLDDT = 91.1%) predicted structure of a Bedadeti protein showing homology with NTP_transf_9 domain-containing protein (UniProt accession ID A0A261RLY6) as indicated in (C). Protein structures were predicted by AlphaFold2 and assessed for structural similarity in Foldseek. Each residue in the sequence is colour-coded based on the model confidence score (pLDDT). The predicted aligned error (PAE) rank box indicates the selected AlphaFold2 model.

Interestingly, in the banana genome, a higher proportion (28 out of 225) of high-confidence structures (pLDDT > 70%) were predicted for MA unique proteins but only a single protein with pLDDT of 77.78% in MB (Supplementary Table 15). Most MB unique proteins reside on scaffolds not assigned to chromosomes and predictions at best fall into the low-confidence (50% < pLDDT < 77.78%) category, possibly representing truncated genes resulting from incomplete assembly. All unique proteins were self-aligned with the predicted structure of MA and MB proteins available in the database. A single MA protein was predicted to be an O-methyltransferase activity (pLDDT = 80.35%, Figure 8A and 8B).

**Figure 8:**
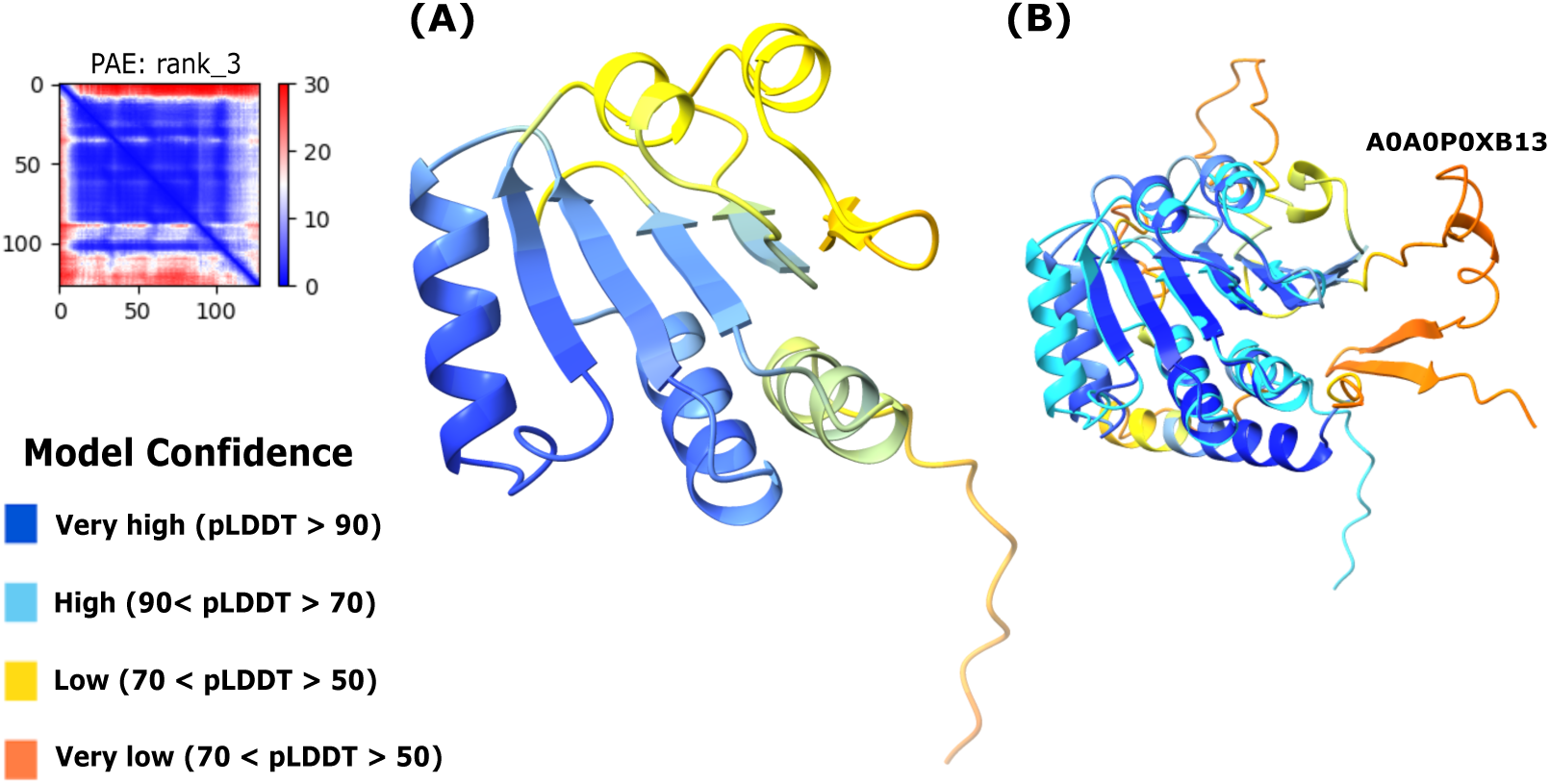
Predicted 3D structure of selected unique proteins in *Musa acuminata* (MA) and their similarity with distantly related proteins. (A) A 127 amino acid residue protein structure (pLDDT = 80.35 %) of an MA unique gene showing disordered N-terminal structure and a relatively low confidence C-terminal structure. (B) A MA unique protein prediction in Figure A (coloured vivid cyan in Figure B) showing structural similarity with O-methyltransferase (UniProt accession ID A0A0C7K9B0). Protein structures were predicted by AlphaFold2, with sequences colour coded with respect to the confidence score (pLDDT) and were assessed for structural similarity in Foldseek. The predicted aligned error (PAE) rank box in the top right of the figure indicates the selected AlphaFold2 model.

### 3.6 Identification of NLR immune receptors

Intracellular nucleotide binding domain leucine rich repeat (NLR) immune receptors are central to plant defence against pathogens, directly or indirectly recognising pathogen effectors to trigger effector-triggered immunity (ETI). NLR gene families exhibit rapid birth-death evolution, with significant copy number variation even among closely related species and cultivars [85, 69, 60, 64]. Given the importance of NLRs in disease resistance and the emerging threat of Xanthomonas wilt (XW) to enset and banana cultivation, we analysed the putative NLR repertoire in enset and compared it with the well-annotated banana genomes.

Using assembled genomic sequences, NLR-Annotator [125] predicted 58 and 64 NLR loci in EV landraces Bedadeti and Mazia, respectively, approximately half the number identified in banana genomes (92 in MB and 127 in MA) (Figure 9A). The relative patterns of NLR class distribution provide informative insights into the enset immune receptor repertoire and are useful for comparing homologous NLRs in draft EV genomes. Our analysis found 127 NLR loci in *M. acuminata* (MA) [9] plus one additional locus that was both annotated as a partial (pseudogene) and a complete (pseudogene), with most loci organised into clusters (2 on Chr3, 1 on Chr7 and 1 on Chr10) (Supplementary Figure 5).

**Figure 9:**
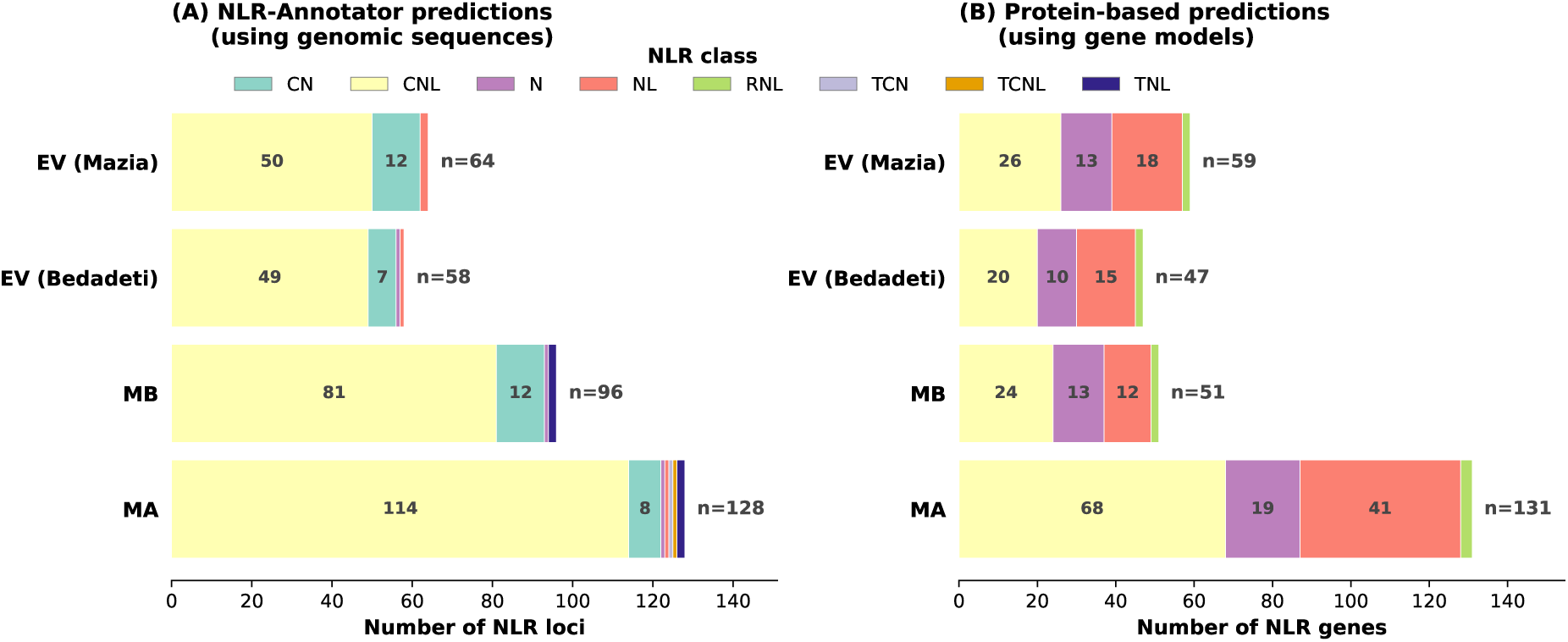
Nucleotide binding domain leucine rich repeat (NLR) immune receptor gene classes in enset landraces and *Musa* species. (A) NLR loci predicted by NLR-Annotator [125] from genomic sequences in *E. ventricosum* (EV) landraces, and *M. acuminata* (MA) and *M. balbisiana* (MB). (B) NLR-encoding genes predicted from annotated protein coding genes in genomes of EV landraces, MA and MB. Numbers within bars indicate counts (>5) per NLR class; total counts shown at bar ends. NLR class abbreviations: CN, CC-NB-ARC (truncated, lacking LRR); CNL, CC-NB-ARC-LRR (complete CC-type NLRs); N, NB-ARC only; NL, NB-ARC-LRR (lacking N-terminal domain—note that CC domain detection varies by CNL subclass (36-76%) [50], so some NL-classified genes may represent CNLs with undetectable CC domains); RNL, RPW8-NB-ARC-LRR; TNL, TIR-NB-ARC-LRR; TCNL, TIR-CC-NB-ARC-LRR; TCN, TIR-CC-NB-ARC. Domain abbreviations: CC, coiled-coil; NB-ARC, Nucleotide Binding site shared with APAF-1, R proteins, and CED-4; LRR, leucine-rich repeat; RPW8, Resistance to Powdery Mildew 8-type domain; TIR, Toll/interleukin-1 receptor. See Supplementary Figure 7 for NLR domain architecture schematic representation.

Cross-species comparisons identified enset-specific NLR loci lacking homologues in both Musa species: 6 in EV landrace Bedadeti and 5 in EV landrace Mazia (Supplementary Table 10). Of these, NLR-Annotator [125] classified two as “complete” with intact reading frames, while the remaining were classified as “complete (pseudogene)” or “partial (pseudogene)”. This could represent either a trace of pseudogenised sequence, an assembly artefact or a potential novel NLR locus. BLAST analysis of these enset-specific NLR loci against nucleotides or proteins from other species revealed that 9 out of 11 showed similarity with other disease resistance genes, reinforcing that these loci are likely potential NLR encoding genes in enset, providing confidence we may have identified novel NLR loci.

Complementary analysis using annotated protein coding genes identified 47 and 59 NLR-encoding genes in EV landrace Bedadeti and Mazia genomes, respectively (Figure 9B). In the Musa genomes, 51 NLRs were predicted in MB protein coding genes compared with 131 in MA. These predictions were generally more conservative than those of NLR-Annotator results (which predicted 58/64 in EV landrace Bedadeti/Mazia and 92/127 in MB/MA) and represent a combination of both variation in the assembly quality and biological differences between the species. It is highly unlikely that the former accounts for the nearly two-fold higher prediction of NLRs in MB by NLR-Annotator. The notably lower number of predicted NLR genes in MB might simply reflect a stringency in the NLR gene prediction pipeline or represent potential NLR encoding genes missed in the MB annotation pipeline.

To assess our predictions, NLR genes were mapped back to their respective genomes. These NLR genes overlapped with the majority of NLR loci predicted by NLR-Annotator: 55 of 58 loci in EV Bedadeti and 61 of 64 loci NLR loci in EV Mazia, along with 83 of 92 loci in MB and 113 of 127 NLR loci in MA, showed correspondence with predicted NLR encoding genes (Supplementary Figures 5 and 7). The remaining 66 loci without gene annotation support were queried against the NCBI non-redundant protein or nucleotide collection database using BLASTX or BLASTN search. Of these, 65 (98.5%) had significant BLAST hits, with 62 (93.9%) showing similarity to characterised disease resistance proteins, including RGA family members (25 loci), RPP13-like proteins (11 loci), generic resistance proteins (10 loci), Pik-2-like proteins (3 loci), CC-NBS-LRR (3 loci), and TIR-NBS-LRR (2 loci). The proportion of loci with disease resistance homology was high across all genomes (MA: 100%; MB: 92.3%; EV Bedadeti: 90.0%; EV Mazia: 88.9%), reflecting the majority of the predicted NLR loci encode potential disease resistance-related proteins.

### 3.7 Phylogenetic analysis of NLRs

To understand the evolutionary relationships among NLRs in enset and banana (MA and MB), we performed phylogenetic analysis using the highly conserved NB-ARC domains. The phylogenetic analysis revealed differences in NLR family expansion between enset and banana (Figure 10). Enset harbours fewer monophyletic NLR clades compared with *Musa* species, with the majority of expanded clades derived from MA. This pattern is consistent with a previous report of NLR gene family expansion in MA and their reduced representation in MB [140]. Interestingly, resistance to EXW exists in MB [102]. Though fewer NLR clades were expanded in EV compared to *Musa* species, the phylogenetic analysis identified multiple clusters of enset specific NLR gene families (highlighted green or light green in Figure 10) that may provide a possible unique reservoir that could be exploited for resistance breeding in enset.

**Figure 10:**
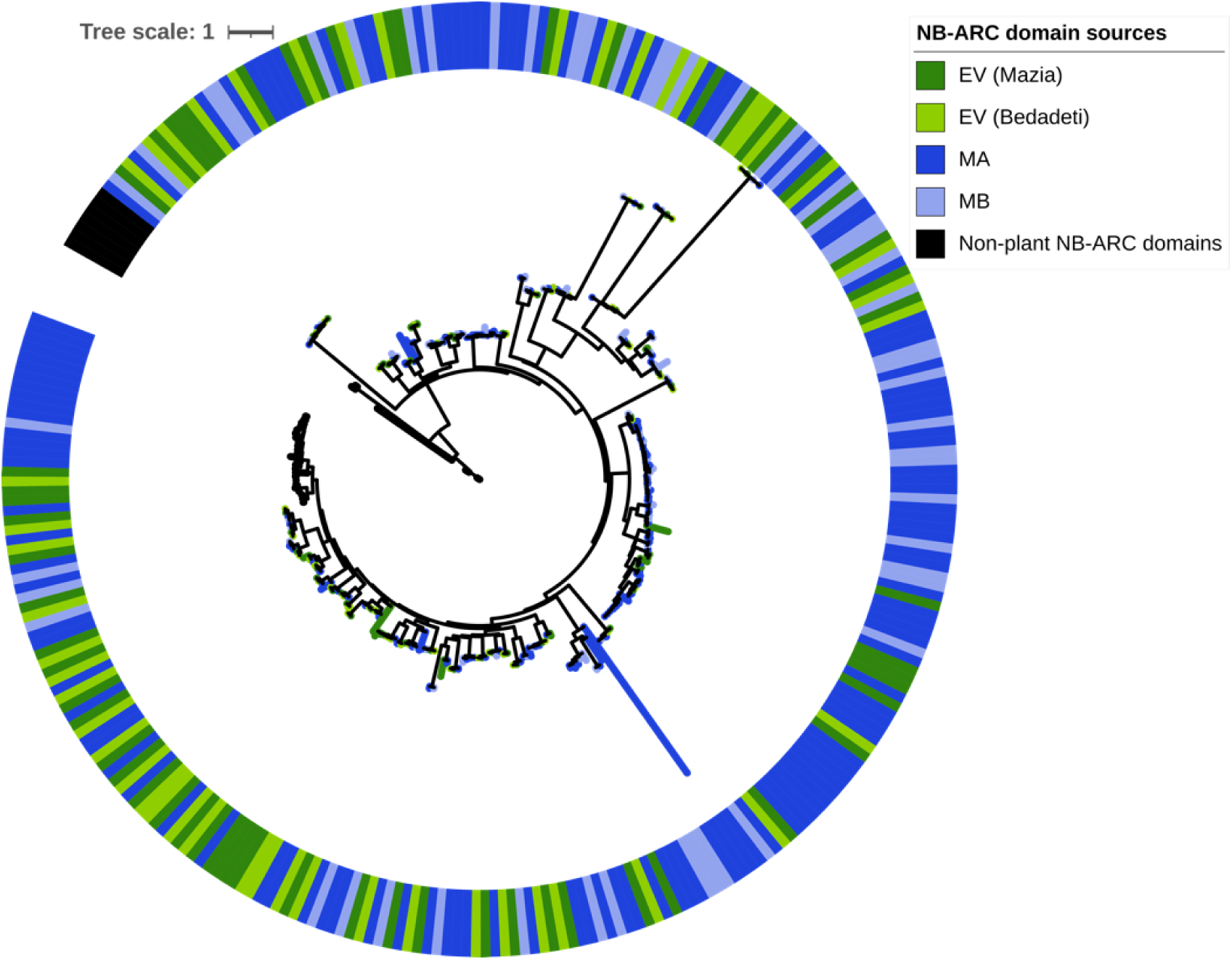
Phylogenetic tree of NLRs from *E. ventricosum* (EV) landraces Mazia and Bedadeti, *M. acuminata* (MA) and *M. balbisiana* (MB). The outer encircled coloured ring represents source of input NB-ARC domain from EV landrace Mazia and Bedadeti, and *Musa* species (MA and MB) as indicated in the legend label. The tree was generated using RAxML (v8.2.12) [124] applying the Maximum Likelihood method based on the JTT model and bootstrap value of 1000 replicates. For outgroups rooting, non-plant NB-ARC containing domains from RefPlantNLR [80] were included and the tree was annotated using iTOL [81].

Consistent with the evolutionary loss of TIR-NLRs (TNLs) in monocots, no genomic loci harbouring either Toll/interleukin-1 receptor (TIR) domain-containing NLR (TNL) or TIR encoding NLR [11] genes were detected in enset. Although NLR-Annotator predicted four loci harbouring TNL-like loci in *M. acuminata*, detailed domain analysis revealed these loci showed no detectable TIR signatures, suggesting these as false positives. These are unlikely to be genuine TN/TNLs as TIR NLRs are typically absent or found in strictly limited numbers in monocots [130].

Among non-canonical NLR classes, we identified 4 and 5 Resistance to Powdery Mildew 8 (RPW8)-domain containing NLRs (RNLs) in enset and *Musa* gene models, respectively. RNLs, also known as “helper” NLRs, are involved in transducing signals from “sensor” NLRs [67], predominantly TNLs, to activate the effector triggered hypersensitive response in plants [79, 27]. Phylogenetic analysis indicated that these “helper” RNLs appear to have recently evolved or expanded paralogue pairs in enset and *Musa* genomes. The conservation of RNLs in enset, despite its reduced overall NLR repertoire, indicates that the core immune signalling machinery remains intact and may provide a foundation for functional disease resistance even with fewer sensor NLRs.

## 4 Discussion

*Ensete ventricosum* (EV), a unique and multipurpose orphan crop, is cultivated in south and southwestern Ethiopia, supporting over 20 million people for both food and non-food purposes. Despite its contribution to food security and resilience, genomic analysis of enset lags behind regionally important crops like banana [9, 140] and other cash crops. Surprisingly, given its immense value to Ethiopia, genomic resources for enset breeding and improvement are virtually non-existent [12, 45, 112]. To address this, we generated a genome sequence for enset landrace Mazia, which is resistant to EXW. This represents a substantial improvement on our previously published genome sequence for landrace Bedadeti and enabled comparisons between enset and two related *Musa* banana genomes.

Using our enset assemblies we undertook comparative analyses with high-quality, annotated genome assemblies of the two major progenitors of edible bananas, *M. acuminata* (MA) and *M. balbisiana* (MB). Despite being phenotypically and phylogenetically related, the genomes of the three species vary considerably. EV shows a marked 10-12% higher abundance of repetitive elements. As enset is clonally cultivated, it is highly likely the stress associated with this could activate distinct TE families through *cis* regulatory elements, epigenetic marks [48], or other mechanisms [33, 88, 114]. Indeed, it is well documented that transposition is enhanced by pathogen infection [156, 5], wounding [114, 95] and environmental stresses [40, 144, 110]. As domesticated enset is propagated vegetatively, in the absence of the sexual recombination seen in pollinated crops, we speculate that TE activation may underlie the phenotypic plasticity and local adaptation evident in today’s myriad enset landraces [101]. Further investigation of the mechanistic role of TEs in shaping the enset genome and its domestication is warranted.

Surprisingly, we found from the EV, Musa comparative analyses, that 25% of the enset genome was not conserved in *M. acuminata* or *M. balbisiana*. Strikingly, these unique enset sequences occupy about 7% of enset’s gene space and likely reflect the biological divergence that occurred during enset and banana evolution and subsequent domestication following their split from their last common ancestor ∼51.9 Mya [65, 39].

At the protein level, we detected a large complement of enset-specific genes we predict contribute to the lifestyle, adaptation and corm-productive qualities underpinning enset’s agronomic traits. These families include proteins with putative roles in DNA integration (consistent with high TE gene content), carbohydrate metabolism (possibly underpinning corm development), and responses to abiotic and biotic stresses such as drought and disease resistance. Rather than producing edible fruit, enset generates a large corm and pseudostems that are rich in starch. These phenotypic characteristics are consistent with the identification of genes in EV that are associated with carbohydrate metabolism. Stress-induced birth of distinct transposable element families has been documented in other crop species [48, 33, 88, 114]. We hypothesise that the enset-specific transposition genes may have driven widespread re-modelling of the enset genome, activated by the extensive physical damage to the apical meristem during clonal cultivation. Future studies warrant comparing the TE landscapes of wild enset as well as domesticated enset.

Identified genes specific to *Musa* species included genes associated with fruit development and flowering, consistent with enset’s lifestyle; enset flowers once after about 8–12 years and rarely produces an edible fruit. Similarly, species-specific gene families associated with resistance and response to biotic and abiotic stresses were identified as unique to *Musa*, consistent with recent studies on *Musa* genomes [9, 140, 30], possibly reflecting different local selection pressures.

We used AlphaFold2 to identify structural homologues of enset’s orphan genes in distantly related species. Surprisingly, among the genes unique to enset, AlphaFold2 predicted a structure with high confidence for only 2% of EV specific genes whereas in *M. acuminata* this was 13%. Among the proteins unique to enset, a relatively high proportion are predicted to be disordered, compared with proteins unique to *Musa* spp. Disordered proteins often function as scaffold proteins [94, 119, 134], are frequently involved in adaptation to stress [98, 135] but are difficult to model. One may speculate that orphan genes in enset and banana that lack detectable homology may contribute to facilitating lineage-specific novel adaptations to changing environmental conditions. Indeed, it has been proposed that orphan genes contribute to lineage specific adaptation [132], tolerance to disease [97, 146] and drought [37, 96], and could thus help facilitate enset’s domestication dynamics.

Enset and *Musa* species also differ in their modes of perenniality and growth morphologies. Cultivated enset has an extended juvenile-to-vegetative growth phase, taking up to 12 or more years. In contrast, *Musa* species often flower within 1.5 years [96]. Enset differs fundamentally from banana in formation of an underground, energy-rich starchy corm, which serves as an important food reserve under drought and stress conditions. Consistent with this, we identified EV specific genes implicated in biological processes associated with allocation of resources/nutrient storage, energy and survival strategies, possibly contributing to prolonging lifespan as in other perennial plant species [49, 120, 56, 6]. In contrast, we identified *Musa* species-specific gene families implicated in metabolism associated with fruit development [140]. In summary, these species-specific sets of genes in EV, MA and MB appear to reflect fundamental phenotypic and lifestyle differences between enset and *Musa* species.

Finally, we extended our analysis to comparative profiling of nucleotide binding domain leucine rich repeat (NLR) family disease resistance (R) gene repertoires. The enset landrace Mazia genome harbours only half (46%–50%) of the *R* gene families encoded in MA genome with a similar NLRs reduction pattern in landrace Bedadeti, and MB. This finding is consistent with previous work reporting that NLR gene family expansion is associated with host-pathogen interactions in MA, and their reduced representation in MB [140]. The reason for low numbers of NLR genes encoded by enset is unclear. Moreover, we also identified multiple enset and banana specific NLR encoding genes. With disease pressure becoming an increasing problem in enset and banana cultivation, particularly EXW and enset landrace Mazia showing high tolerance to EXW, this NLR gene repertoire provides a potentially unique resource that could be exploited for resistance breeding.

Furthermore, we identified differences between genomic and protein-based NLR predictions in counts and NLR class detection, which is expected and biologically meaningful rather than a methodological artefact in our analysis. Genomic approaches detect all sequences containing NLR-associated motifs [125], including pseudogenes with frameshift mutations or premature stop codons, unannotated genes, and partial sequences from assembly fragmentation. In contrast, protein-based approaches require complete open reading frames and depend on gene annotation quality. Interestingly, a recent study reported variable coiled-coil (CC)-domain detection rates (36-76%) of CC-NLR [50]. This biological variation in CC domain detectability appears to have functional significance [51] but has important implications for NLR classification in a way that many functional CNLs may be misclassified as “NL” (lacking N-terminal domain) simply because their CC domains fall below detection thresholds, rather than representing genuinely truncated proteins. Interestingly, in the wheat genome, only 46% of genomic NLR loci corresponded to validated protein-coding genes with intact ORFs [125], a ratio similar to our observations in MB. The discrepancy is particularly pronounced in MB, where NLR-Annotator predicted nearly twice as many loci as the protein-based approach which likely reflect both incomplete gene annotation in NLR clusters and the presence of recently pseudogenised NLR loci that may retain detectable sequence motifs. Given the draft nature of our enset assemblies the reported NLR counts likely represent minimum estimates; chromosome-scale assemblies may reveal additional NLR loci currently fragmented across contigs or located in assembly gaps. Notably, even telomere-to-telomere sequencing of Cavendish banana revealed 123-140 NLR loci across its three haploid genomes [60], compared to 127 in the MA reference, illustrating that substantial NLR copy number variation exists even within species and among haplotypes of cultivars.

This study was constrained by differences in the genome sequencing platforms on which the data were generated, subsequent differences in assembly metrics, and the reliance on putatively inferred gene functions. Our NLR analysis was entirely computational, based on sequence similarity and domain prediction. The comparative patterns observed between species nonetheless provide initial insights into enset immune receptor architecture. Consequently, more comprehensive comparisons of the enset and *Musa* species genomes, along with their domestication signatures, and the validation of putatively identified gene functions were not possible. However, with the advancement of telomere-to-telomere scale genome sequencing, future research can advance our comprehension of enset domestication, particularly the differences between wild and cultivated enset. This knowledge can then be applied to enhance our understanding of common biotic and abiotic threats and support the development of molecular-based enset trait selection strategies.

## 5 Data Availability

Raw sequenced reads have been uploaded to the NCBI Sequence Read Archive under accession numbers SRR27722736 for short stLFR reads and SRR27722998, SRR27722999, and SRR27723000 for RNA-seq reads. The assembled genome is available under accession number GCA_052324855.1. Annotations and other files have been submitted to Zenodo [99].

## 5.1 List of abbreviations

EV: Ensete ventricosum
MA: Musa acuminata
MB: Musa balbisiana
NLR: Nucleotide-binding Leucine-rich Repeat
NB-ARC: Nucleotide-Binding adaptor shared by APAF-1, R proteins, and CED-4
LRR: Leucine-Rich Repeat
CC: Coiled-Coil
TIR: Toll/Interleukin-1 Receptor
RNL: RPW8-NB-LRR (helper NLR)
CNL: CC-NB-LRR
TNL: TIR-NB-LRR
EXW: Enset Xanthomonas Wilt
BXW: Banana Xanthomonas Wilt
GO: Gene Ontology
TE: Transposable Element
LTR: Long Terminal Repeat
CDS: Coding Sequence
ORF: Open Reading Frame
stLFR: single-tube Long Fragment Read
BUSCO: Benchmarking Universal Single-Copy Orthologs
pLDDT: predicted Local Distance Difference Test

## 5.2 Competing Interests

The authors declare that they have no competing interests.

## 5.3 Funding

This study was supported by the University of Warwick Chancellor’s International Fellowship (to SM), BGI Health (HK) Company Limited, BBSRC GCRF Accelerator Fund (BB/M017982/1) (to MG), Royal Society Challenge Grants (CHG\R1\170082) (to VN), and UKRI/BBSRC: BB/M017982/1 Warwick Integrative Synthetic Biology Centre funding (to AD-F). SKS acknowledges 10KP project. The funders had no role in study design, data collection and analysis, decision to publish, or preparation of the manuscript.

## 5.4 Authors’ Contributions

SM, DJS and MG analysed and interpreted the data. SM wrote the original manuscript draft. PP and AD-F contributed to methodology and investigation. SKS, AVD, GM, ZY, KT and VN provided resources. DJS and MG supervised the project.

## 6 Acknowledgements

The authors thank Southern Agricultural Research Institute for providing samples as well as necessary services and facilities during sample collection, Bio and Emerging Technology Institute (BETin) for facilitating DNA extraction and Ethiopian Biodiversity Institute (EBI) for facilitating sample export for sequencing. We used computing resources provided by the University of Warwick Scientific Computing Research Technology Platform (SCRTP).

---

**Supplementary Figure 1.**
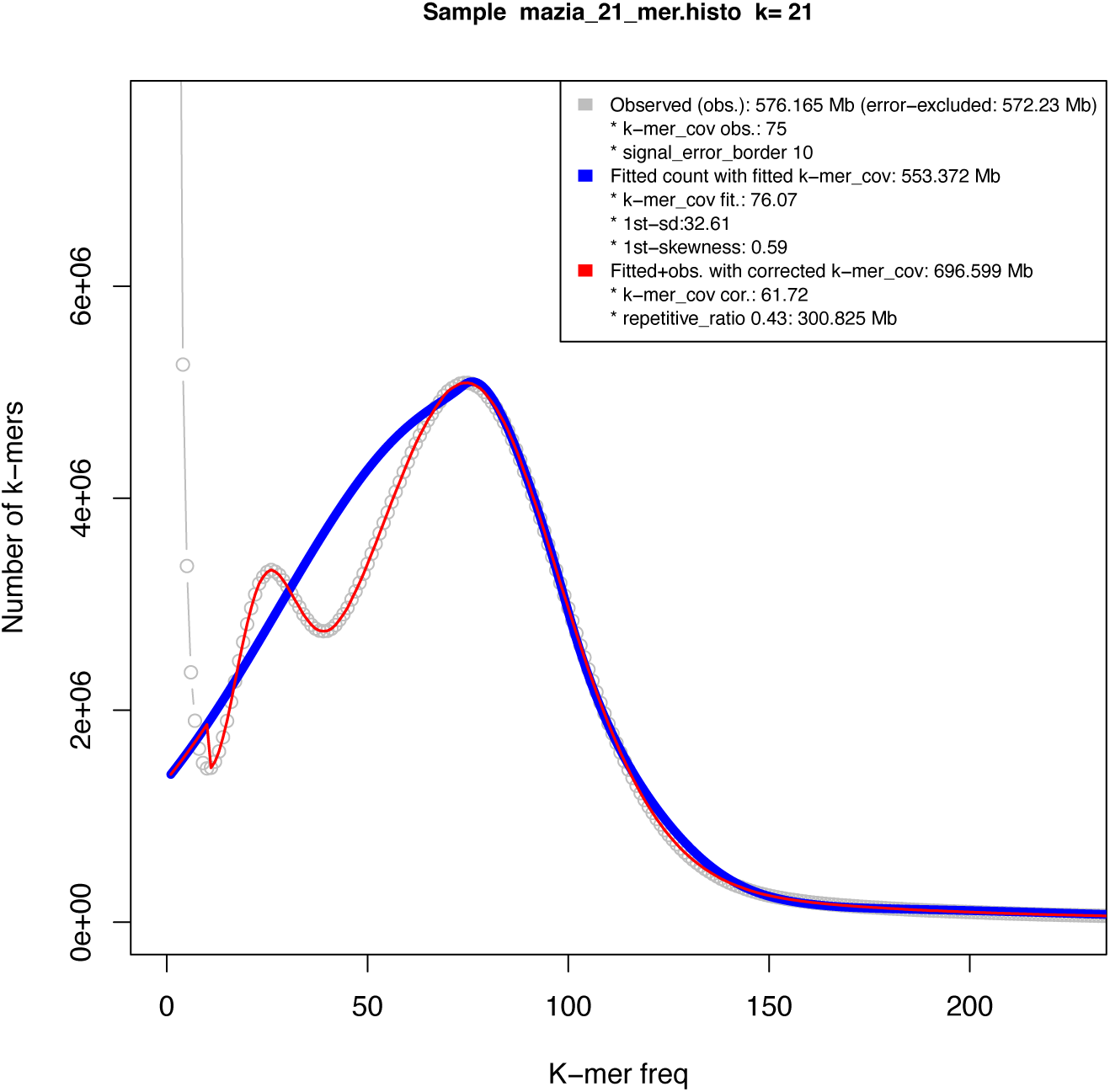
Estimated genome size of enset (*Ensete ventricosum*) landrace Mazia as inferred by findGSE.

**Supplementary Figure 2.**
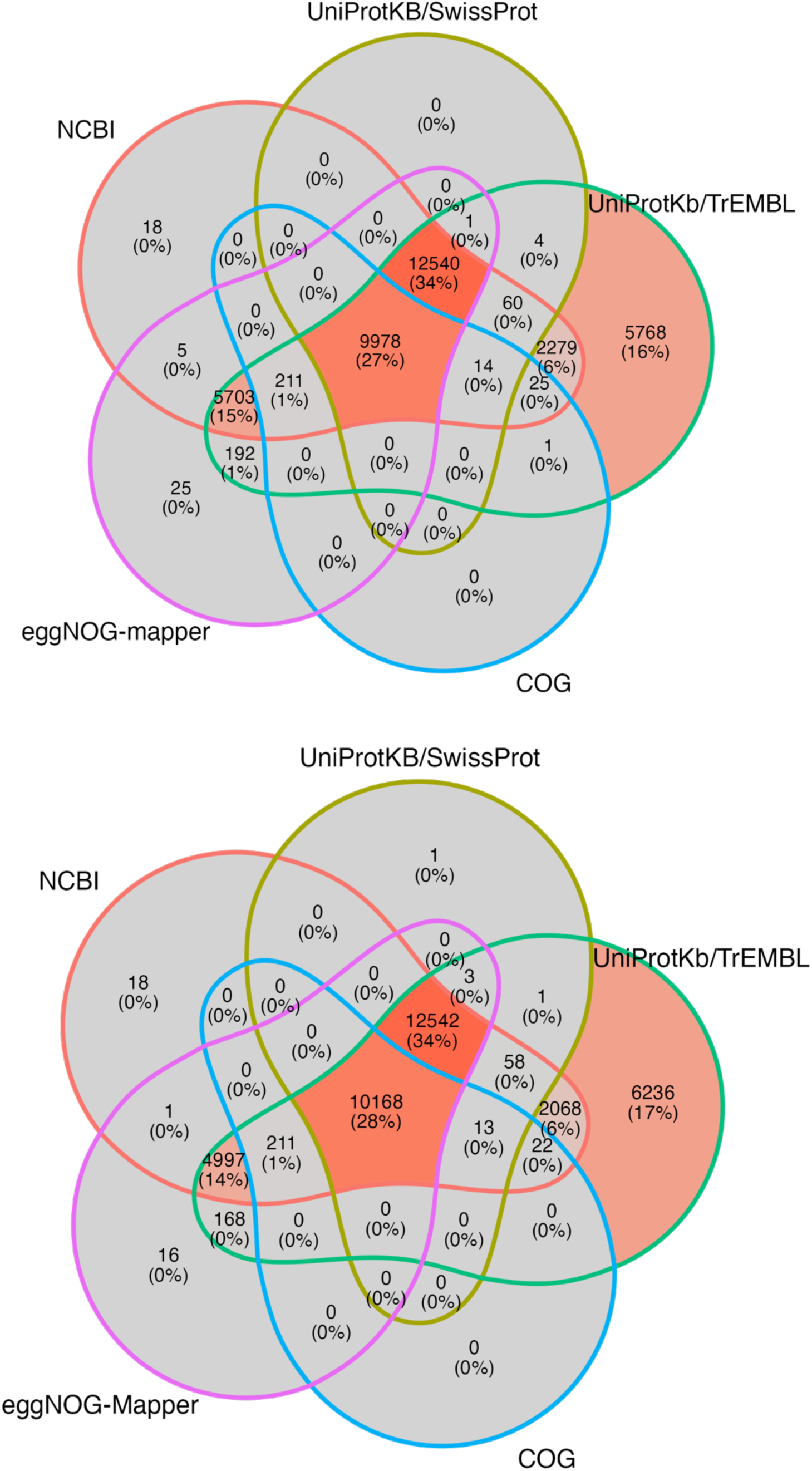
Functional annotation of protein coding genes for enset (*Ensete ventricosum*) landrace Mazia (top) and Bedadeti (bottom)

**Supplementary Figure 3.**
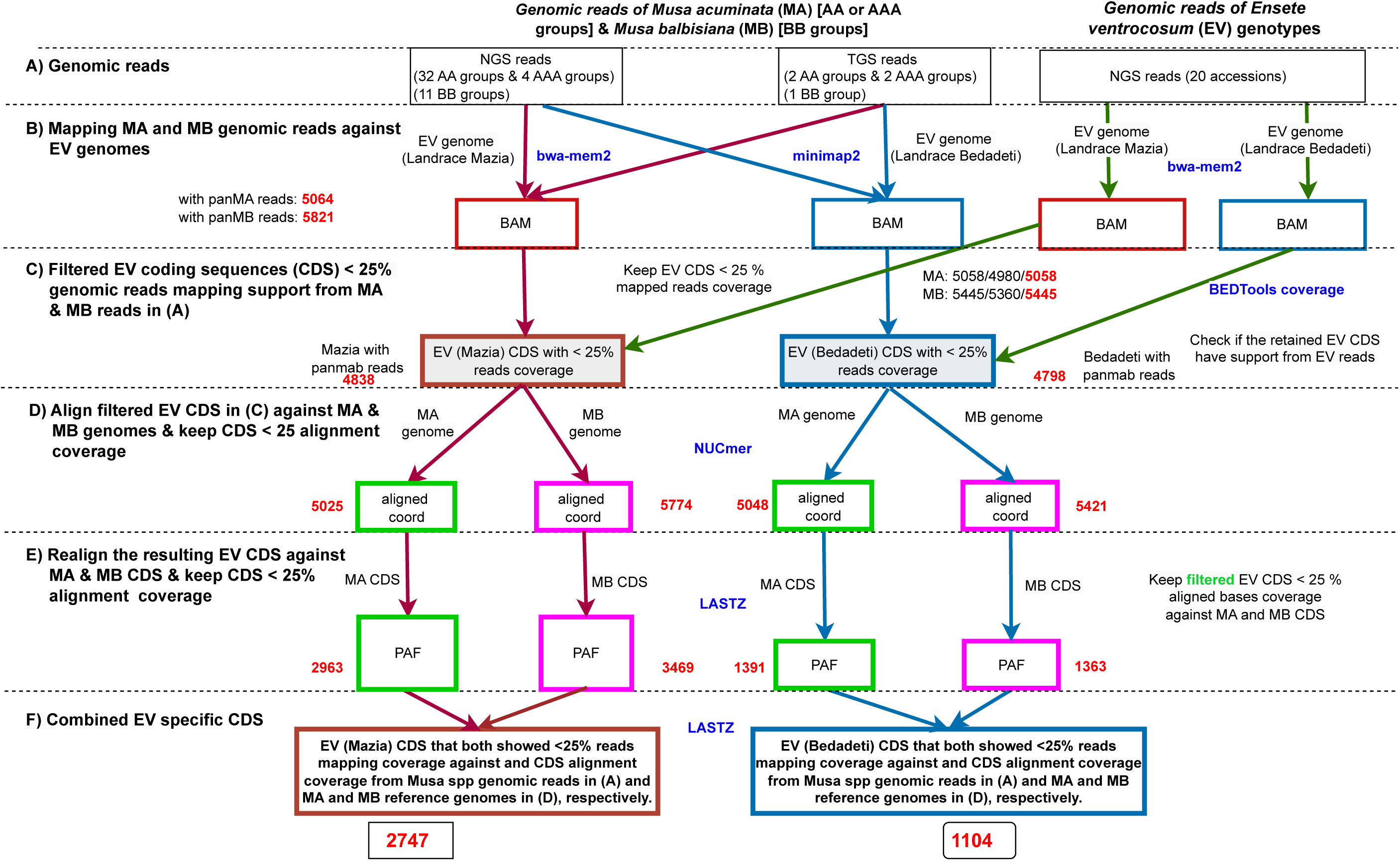
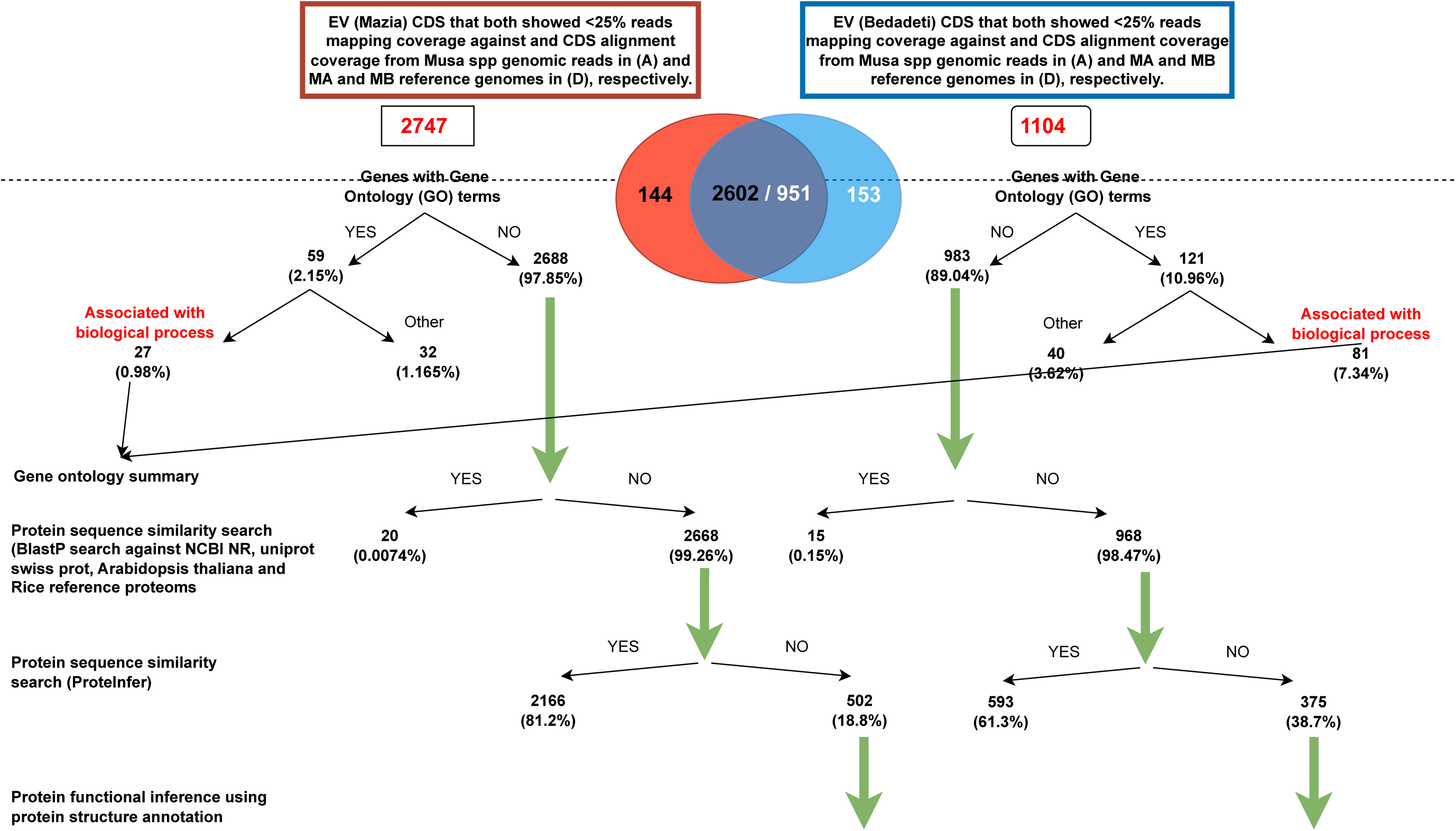
Data workflow to identify enset (*E. ventricosum*) specific protein coding genes using genomic reads mapping and coding sequences alignment.

**Supplementary Figure 4.**
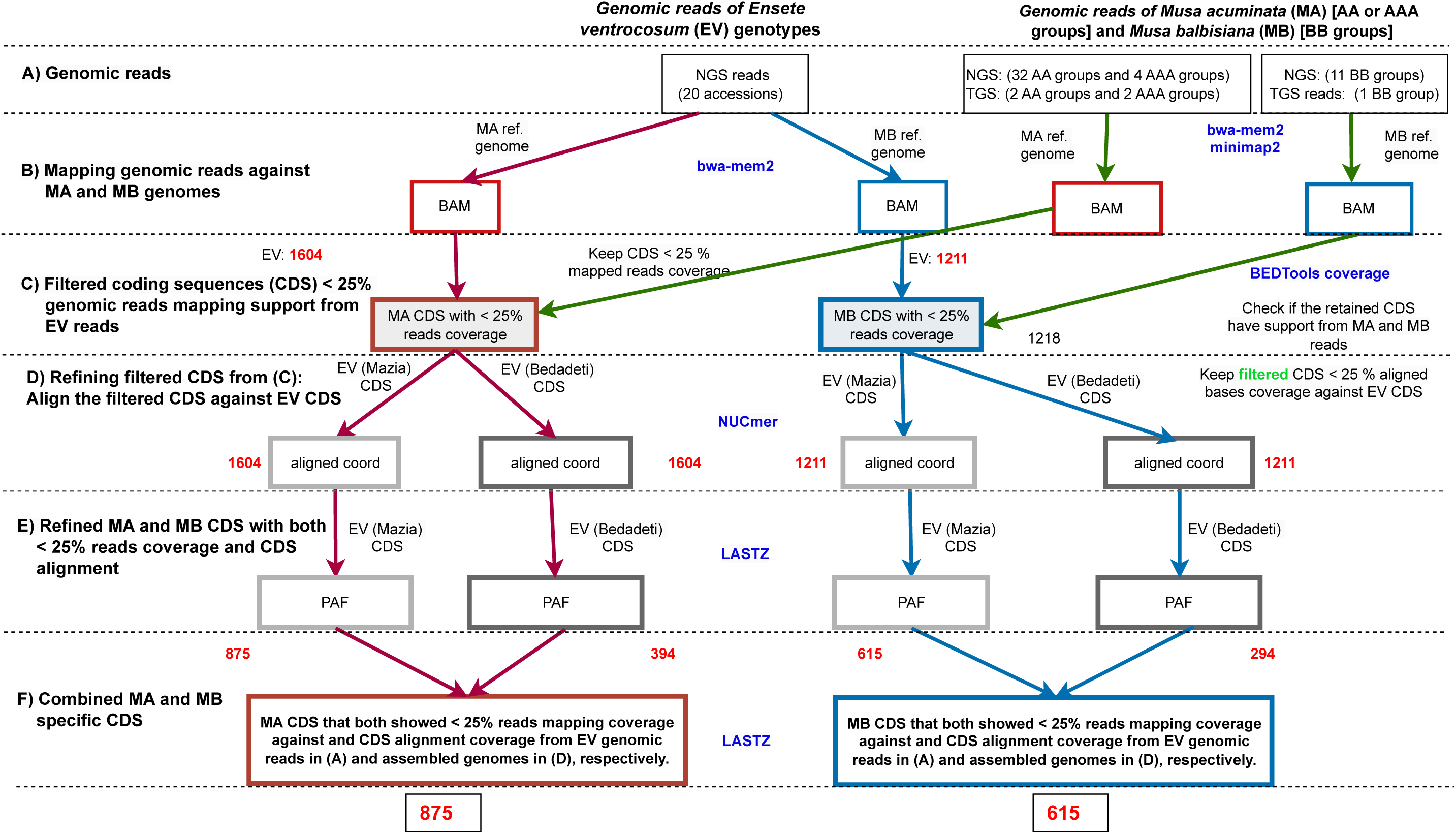
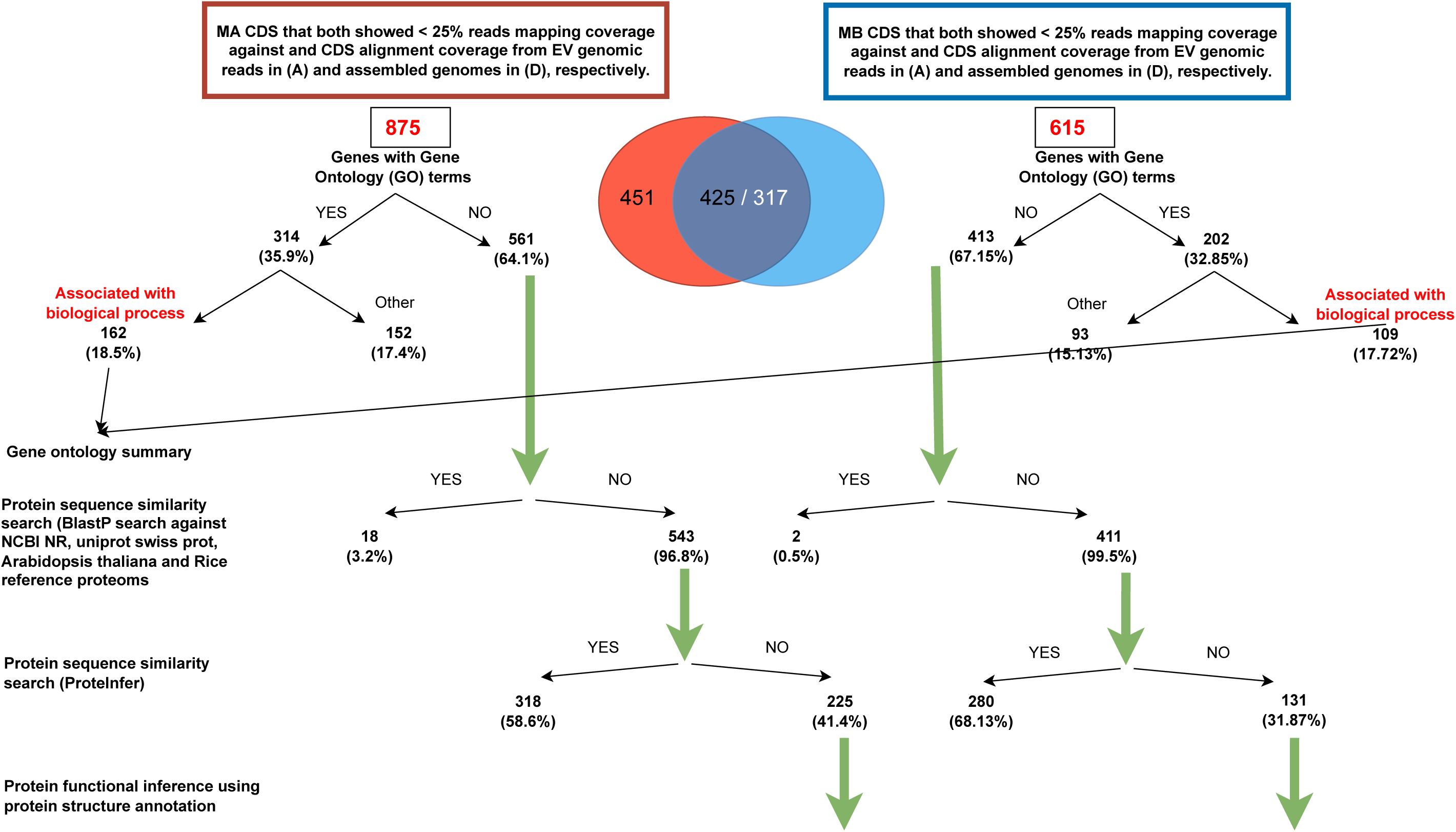
Data workflow to identify *M. acuminata* (MA) and *M. balbisiana* (MB) specific protein coding genes using genomic reads mapping and coding sequences alignment

**Supplementary Figure 5.**
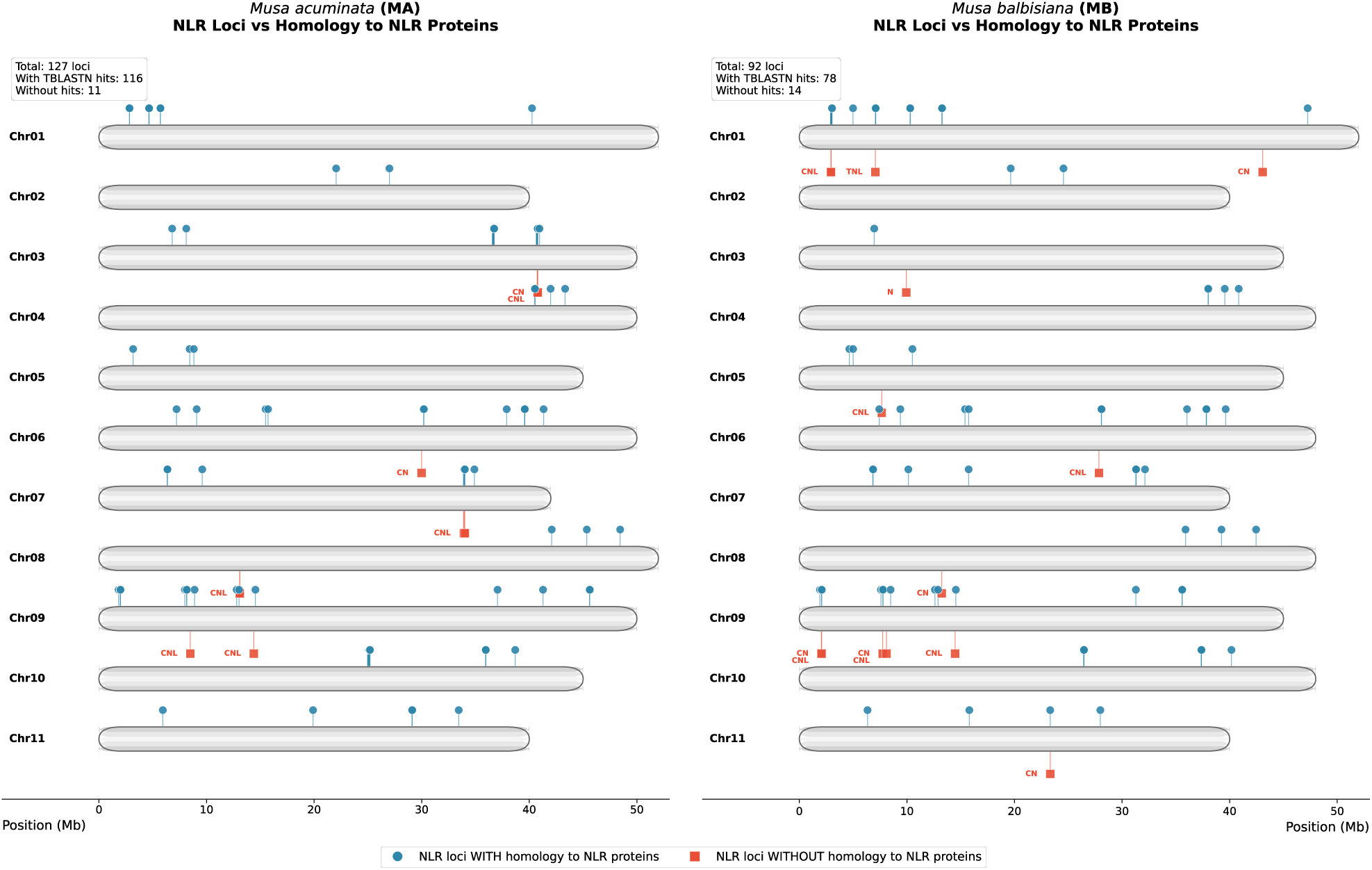
Predicted NLR loci distribution in *Musa acuminata* and *Musa balbisiana* genomes.

**Supplementary Figure 6.**
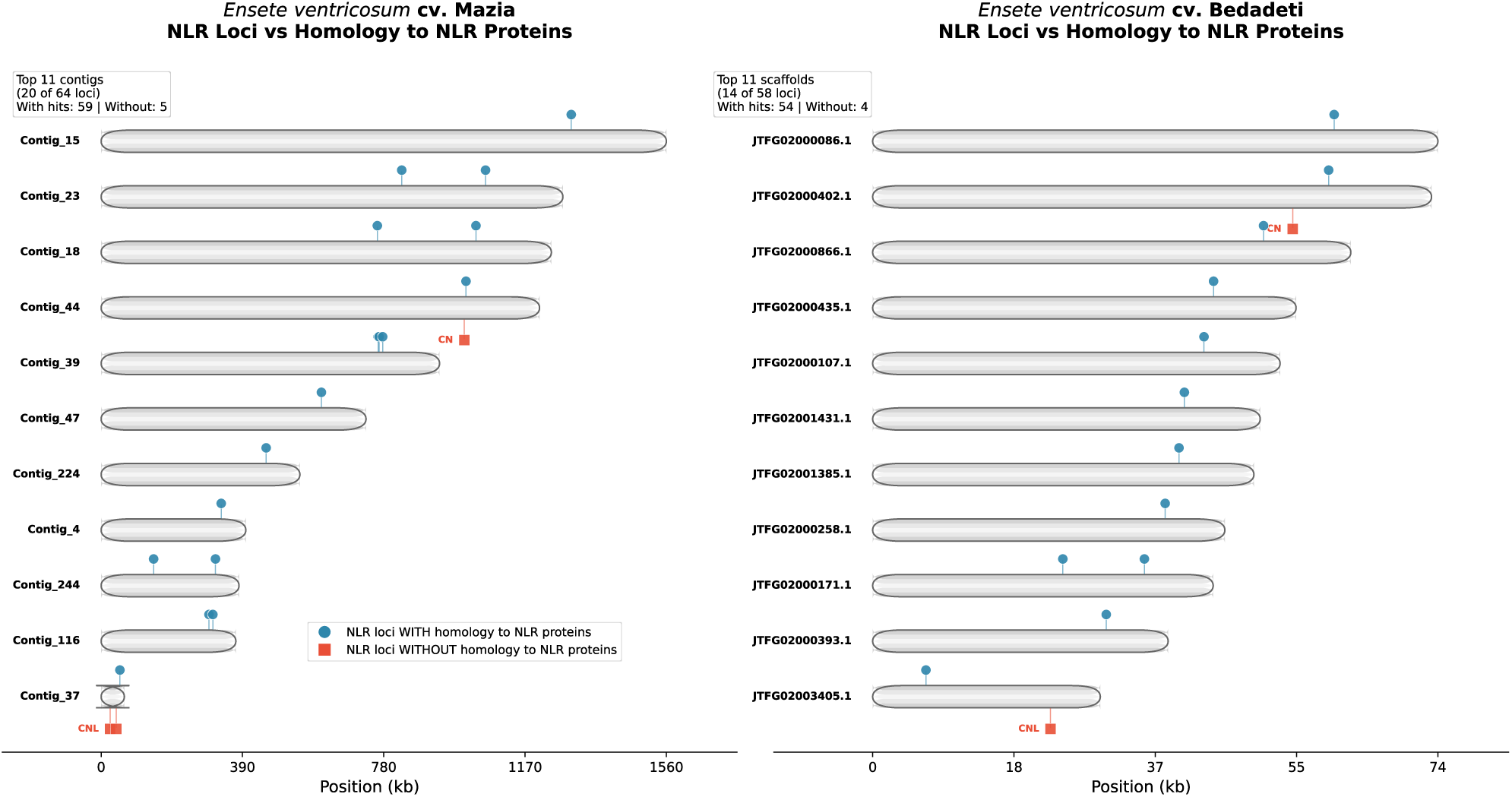
Predicted NLR loci distribution in selected contig of enset (*Ensete ventricosum*) landrace Mazia and Bedadeti selected genomes.

**Supplementary Figure 7.**
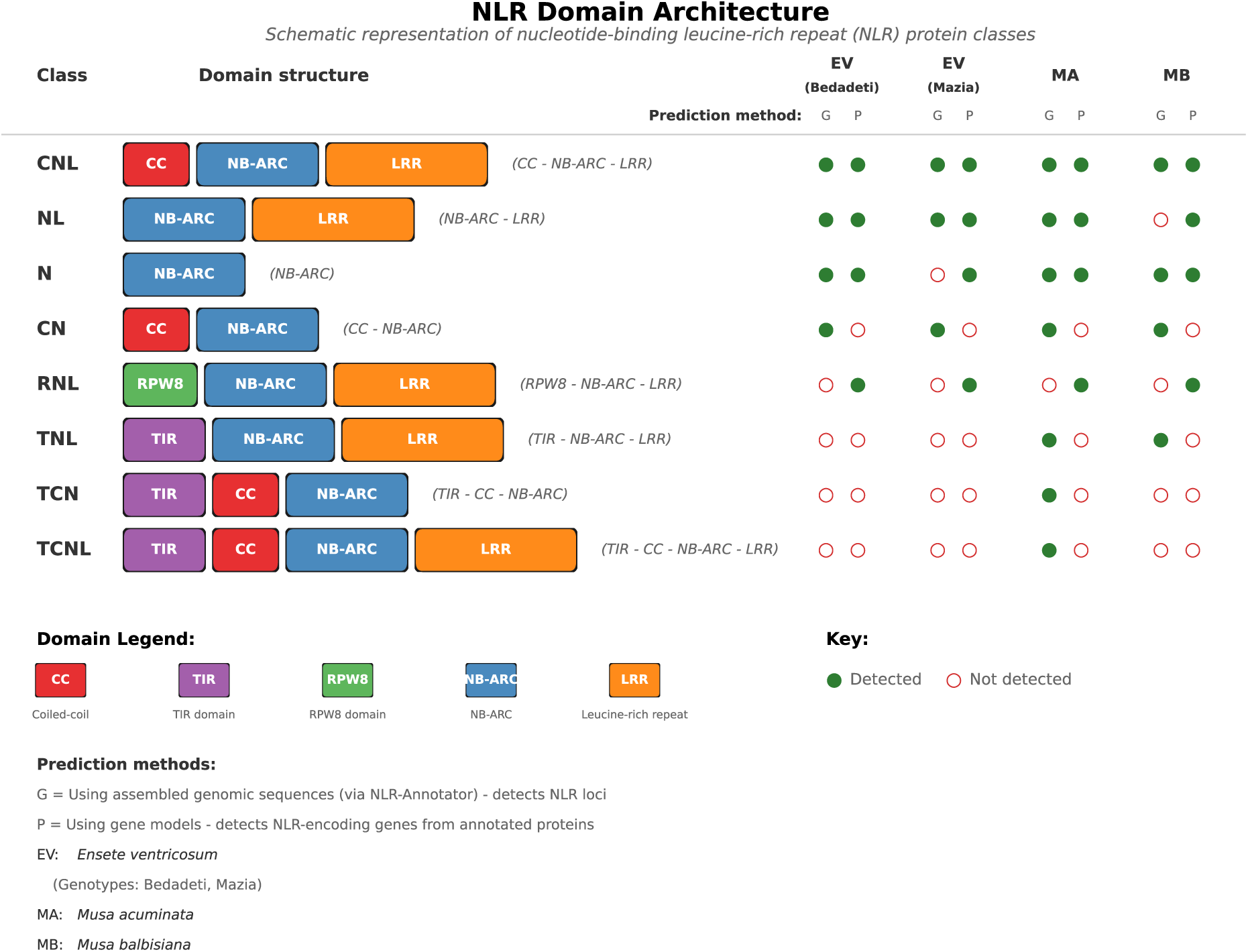
Summary of NLR domain architechture prediction from genomic sequence and protein coding genes in enset (*Ensete ventricosum*) landrace Mazia and Bedadeti, Musa species (*Musa acuminata* and *Musa balbisiana*)

## Notes

https://zenodo.org/records/18292608

### Competing Interest Statement

The authors have declared no competing interest.

## References

[1] A. Abraham, S. Winter, K.R. Richert-Pöggeler, and W. Menzel. Molecular characterization of a new badnavirus associated with streak symptoms on enset (*Ensete ventricosum*, musaceae). Journal of Phytopathology, 166(7-8):565–571, 2018.

[2] T. Addis, F. Azerefegne, G. Blomme, and K. Kanaujia. Biology of the enset root mealybug *Cataenococcus ensete* and its geographical distribution in southern ethiopia. Journal of Applied Biosciences, 8(1):251–260, 2008.

[3] S.A. Aleksander, J. Balhoff, S. Carbon, J.M. Cherry, H.J. Drabkin, D. Ebert, et al. The gene ontology knowledgebase in 2023. GENETICS, 224(1), 2023.

[4] S. Andrews. Fastqc: a quality control tool for high throughput sequence data. http://www.bioinformatics.babraham.ac.uk/projects/fastqc, 2010.

[5] K.I. Ansari, S. Walter, J.M. Brennan, M. Lemmens, S. Kessans, A. McGahern, et al. Retrotransposon and gene activation in wheat in response to mycotoxigenic and non-mycotoxigenic-associated fusarium stress. Theoretical and Applied Genetics, 114(5):927–937, 2007.

[6] Y. Aoyagi Blue, J. Kusumi, and A. Satake. Copy number analyses of dna repair genes reveal the role of poly(adp-ribose) polymerase (parp) in tree longevity. iScience, 24(7):102779, 2021.

[7] I. Barrio-Hernandez, J. Yeo, J. Jänes, M. Mirdita, C.L.M. Gilchrist, T. Wein, et al. Clustering predicted structures at the scale of the known protein universe. Nature, 622(7983):637–645, 2023.

[8] P. Bayer. annotateclasses.py. https://gist.github.com/philippbayer/0052f5ad56121cd2252a1c5b90154ed1, 2022.

[9] C. Belser, F.-C. Baurens, B. Noel, G. Martin, C. Cruaud, B. Istace, et al. Telomere-to-telomere gapless chromosomes of banana using nanopore sequencing. Communications Biology, 4(1), 2021.

[10] G. Benson. Tandem repeats finder: a program to analyze dna sequences. Nucleic Acids Research, 27(2):573–580, 1999.

[11] M. Bernoux, T. Ve, S. Williams, C. Warren, D. Hatters, E. Valkov, et al. Structural and functional analysis of a plant resistance protein tir domain reveals interfaces for self-association, signaling, and autoregulation. Cell Host & Microbe, 9(3):200–211, 2011.

[12] J.A. Bhat, D. Yu, A. Bohra, S.A. Ganie, and R.K. Varshney. Features and applications of haplotypes in crop breeding. Communications Biology, 4(1):1266, 2021.

[13] G. Blomme, M. Dita, K.S. Jacobsen, L. Perez Vicente, A. Molina, W. Ocimati, et al. Bacterial diseases of bananas and enset: Current state of knowledge and integrated approaches toward sustainable management. Frontiers in Plant Science, 8:1290, 2017.

[14] M. Blum, H.-Y. Chang, S. Chuguransky, T. Grego, S. Kandasaamy, A. Mitchell, et al. The interpro protein families and domains database: 20 years on. Nucleic Acids Research, 49(D1):D344–D354, 2021.

[15] B. Boeckmann, A. Bairoch, R. Apweiler, M.C. Blatter, A. Estreicher, E. Gasteiger, et al. The swiss-prot protein knowledgebase and its supplement trembl in 2003. Nucleic Acids Research, 31(1):365–370, 2003.

[16] D. Bolser, D.M. Staines, E. Pritchard, and P. Kersey. Ensembl plants: Integrating tools for visualizing, mining, and analyzing plant genomics data. In Plant Bioinformatics, volume 1374, pages 115–140. Springer New York, 2016.

[17] J.S. Borrell, M.K. Biswas, M. Goodwin, G. Blomme, T. Schwarzacher, J.S.P. Heslop-Harrison, et al. Enset in ethiopia: a poorly characterized but resilient starch staple. Annals of Botany, pages 747–766, 2019.

[18] S.A. Brandt, A. Spring, C. Hiebsch, J.T. McCabe, E. Tabogie, G. Wolde-Michael, et al. The “Tree against hunger”: Enset based agricultural systems in Ethiopia. 1997.

[19] Broad Institute. Picard toolkit. https://broadinstitute.github.io/picard/, 2019.

[20] T. Brůna, K.J. Hoff, A. Lomsadze, M. Stanke, and M. Borodovsky. Braker2: automatic eukaryotic genome annotation with genemark-ep+ and augustus supported by a protein database. NAR Genomics and Bioinformatics, 3(1):lqaa108, 2021.

[21] B. Buchfink, K. Reuter, and H.G. Drost. Sensitive protein alignments at tree-of-life scale using diamond. Nature Methods, 18(4):366–368, 2021.

[22] B. Buchfink, C. Xie, and D.H. Huson. Fast and sensitive protein alignment using diamond. Nature Methods, 12(1):59–60, 2015.

[23] M. Busche, B. Pucker, P. Viehöver, B. Weisshaar, and R. Stracke. Genome sequencing of musa acuminata dwarf cavendish reveals a duplication of a large segment of chromosome 2. G3 (Bethesda), 10(1):37–42, 2020.

[24] M.S. Campbell, C. Holt, B. Moore, and M. Yandell. Genome annotation and curation using maker and maker-p. Current Protocols in Bioinformatics, 48, 2014.

[25] C.P. Cantalapiedra, A. Hernández-Plaza, I. Letunic, P. Bork, and J. Huerta-Cepas. eggnog-mapper v2: Functional annotation, orthology assignments, and domain prediction at the metagenomic scale. Molecular Biology and Evolution, 38(12):5825–5829, 2021.

[26] S. Capella-Gutiérrez, J.M. Silla-Martínez, and T. Gabadón. trimal: a tool for automated alignment trimming in large-scale phylogenetic analyses. Bioinformatics, 25(15):1972–1973, 2009.

[27] B. Castel, P.M. Ngou, V. Cevik, A. Redkar, D.S. Kim, Y. Yang, et al. Diverse nlr immune receptors activate defence via the rpw8-nlr nrg1. New Phytologist, 222(2):966–980, 2019.

[28] P.P. Chan, B.Y. Lin, A.J. Mak, and T.M. Lowe. trnascan-se 2.0: Improved detection and functional classification of transfer rna genes. Nucleic Acids Research, 49:9077–9096, 2021.

[29] T. Daba and M. Shigeta. Enset (*Ensete ventricosum*) production in ethiopia: Its nutritional and socio-cultural values. Agriculture and Food Sciences Research, 2016.

[30] A. D’Hont, F. Denoeud, J.-M. Aury, F.-C. Baurens, F. Carreel, O. Garsmeur, et al. The banana (*Musa acuminata*) genome and the evolution of monocotyledonous plants. Nature, 488:213–217, 2012.

[31] C. Diesh, G.J. Stevens, P. Xie, T. De Jesus Martinez, E.A. Hershberg, A. Leung, et al. Jbrowse 2: a modular genome browser with views of synteny and structural variation. Genome Biology, 24(1):74, 2023.

[32] G. Droc, G. Martin, V. Guignon, M. Summo, G. Sempéré, E. Durant, et al. The banana genome hub: a community database for genomics in the musaceae. Horticulture Research, 9:uhac221, 2022.

[33] M.J. Dubin, P. Zhang, D. Meng, M.-S. Remigereau, E.J. Osborne, F. Paolo Casale, et al. Dna methylation in arabidopsis has a genetic basis and shows evidence of local adaptation. eLife, 4, 2015.

[34] B. Dujon. The yeast genome project: what did we learn? Trends in Genetics, 12(7):263–270, 1996.

[35] R.C. Edgar and E.W. Myers. Piler: identification and classification of genomic repeats. Bioinformatics, 21:i151–i158, 2005.

[36] K. Eilbeck, B. Moore, C. Holt, and M. Yandell. Quantitative measures for the management and comparison of annotated genomes. BMC Bioinformatics, 10(1):67, 2009.

[37] A.Z. Fakhar, J. Liu, K.M. Pajerowska-Mukhtar, and M.S. Mukhtar. The lost and found: Unraveling the functions of orphan genes. Journal of Developmental Biology, 11(2), 2023.

[38] J.M. Flynn, R. Hubley, C. Goubert, J. Rosen, A.G. Clark, C. Feschotte, et al. Repeatmod-eler2 for automated genomic discovery of transposable element families. Proceedings of the National Academy of Sciences USA, 117(17):9451–9457, 2020.

[39] N. Fu, M. Ji, M. Rouard, H.F. Yan, and X.J. Ge. Comparative plastome analysis of musaceae and new insights into phylogenetic relationships. BMC Genomics, 23(1):223, 2022.

[40] K. Fujino, S.N. Hashida, T. Ogawa, T. Natsume, T. Uchiyama, T. Mikami, et al. Temperature controls nuclear import of tam3 transposase in antirrhinum. The Plant Journal, 65(1):146–155, 2011.

[41] F. García-Alcalde, K. Okonechnikov, J. Carbonell, L.M. Cruz, S. Götz, S. Tarazona, et al. Qualimap: evaluating next-generation sequencing alignment data. Bioinformatics, 28(20):2678–2679, 2012.

[42] S.L. Gebre, A. Woldeyohannes, K. Getahun, and A. Regassa. Determinants of the spatial distribution of enset (*Ensete ventricosum* welw. cheesman) wilt disease: Evidence from yem special district, southern ethiopia. Cogent Food & Agriculture, 7(1):1889789, 2021.

[43] B. Gel and E. Serra. karyoploter: an r/bioconductor package to plot customizable genomes displaying arbitrary data. Bioinformatics, 33(19):3088–3090, 2017.

[44] B. Gemeda, G. Tesfaye, A. Simachew, A. Wang, A. Mekonnen, A. Guadie, et al. Responses of common enset (*Ensete ventricosum*) varieties in ethiopia to xanthomonas and virulence of xanthomonas strains. Biocatalysis and Agricultural Biotechnology, 53:102872, 2023.

[45] M. Goddard. Genomic selection: prediction of accuracy and maximisation of long term response. In Interbull Bulletin, volume 136, pages 245–257, 2009.

[46] O. Gotoh, M. Morita, and D.R. Nelson. Assessment and refinement of eukaryotic gene structure prediction with gene-structure-aware multiple protein sequence alignment. BMC Bioinformatics, 15(189), 2014.

[47] M.G. Grabherr, B.J. Haas, M. Yassour, J.Z. Levin, D.A. Thompson, I. Amit, et al. Full-length transcriptome assembly from rna-seq data without a reference genome. Nature Biotechnology, 29(7):644–652, 2011.

[48] M.A. Grandbastien. Ltr retrotransposons, handy hitchhikers of plant regulation and stress response. Biochimica et Biophysica Acta, 1849(4):403–416, 2015.

[49] Å. Grimberg, I. Lager, N.R. Street, K.M. Robinson, S. Marttila, N. Mähler, et al. Storage lipid accumulation is controlled by photoperiodic signal acting via regulators of growth cessation and dormancy in hybrid aspen. New Phytologist, 219(2):619–630, 2018.

[50] B.C. Guo, Y.R. Zhang, Z.G. Liu, X.C. Li, Z. Yu, B.Y. Ping, et al. Deciphering plant nlr genomic evolution: Synteny-informed classification unveils insights into tnl gene loss. Molecular Biology and Evolution, 42(2), 2025.

[51] G. Guo, H. Zhao, K. Bai, Q. Wu, L. Dong, L. Lu, et al. An activated wheat cc_*G*10_-nlr immune receptor forms an octameric resistosome. bioRxiv preprint, 2025.

[52] M.F. Hamdan, C.K.S. Karlson, E.Y. Teoh, S.E. Lau, and B.C. Tan. Genome editing for sustainable crop improvement and mitigation of biotic and abiotic stresses. Plants (Basel), 11(19), 2022.

[53] Y. Han and S.R. Wessler. Mite-hunter: a program for discovering miniature inverted-repeat transposable elements from genomic sequences. Nucleic Acids Research, 38:e199, 2010.

[54] R.S. Harris. Improved pairwise alignment of genomic DNA. PhD thesis, The Pennsylvania State University, 2007.

[55] J. Harrison, K. Moore, K. Paszkiewicz, T. Jones, M. Grant, D. Ambacheew, et al. A draft genome sequence for *Ensete ventricosum*, the drought-tolerant “tree against hunger”. Agronomy, 4:13–33, 2014.

[56] D. Herrera-Ramírez, C.A. Sierra, C. Römermann, J. Muhr, S. Trumbore, D. Silvério, et al. Starch and lipid storage strategies in tropical trees relate to growth and mortality. New Phytologist, 230(1):139–154, 2021.

[57] E.A. Hildebrand. A tale of two tuber crops: How attributes of enset and yams may have shaped prehistoric human-plant interactions in southwest ethiopia. In Rethinking Agriculture: Archaeological and Ethnoarchaeological Perspectives, volume 3, pages 273–298. Taylor & Francis group, New York, 2007.

[58] HMMER. Biosequence analysis using profile hidden markov models. http://hmmer.org/, 2024.

[59] K.J. Hoff and M. Stanke. Predicting genes in single genomes with augustus. Current Protocols in Bioinformatics, 65(1):e57, 2019.

[60] H.-R. Huang, X. Liu, R. Arshad, X. Wang, W.-M. Li, Y. Zhou, et al. Telomere-to-telomere haplotype-resolved reference genome reveals subgenome divergence and disease resistance in triploid cavendish banana. Horticulture Research, 10(9), 2023.

[61] R. Hubley, R.D. Finn, J. Clements, S.R. Eddy, T.A. Jones, W. Bao, et al. The dfam database of repetitive dna families. Nucleic Acids Research, 44(D1):D81–D89, 2016.

[62] A.M. Hulse-Kemp, S. Maheshwari, K. Stoffel, T.A. Hill, D. Jaffe, S.R. Williams, et al. Reference quality assembly of the 3.5-gb genome of *Capsicum annuum* from a single linked-read library. Horticulture Research, 5(4):1–13, 2018.

[63] H. Iwata and O. Gotoh. Benchmarking spliced alignment programs including spaln2, an extended version of spaln that incorporates additional species-specific features. Nucleic Acids Research, 40:e161, 2012.

[64] F. Jacob, S. Vernaldi, and T. Maekawa. Evolution and conservation of plant nlr functions. Frontiers in Immunology, 4(297), 2013.

[65] S.B. Janssens, F. Vandelook, E. De Langhe, B. Verstraete, E. Smets, I. Vandenhouwe, et al. Evolutionary dynamics and biogeography of musaceae reveal a correlation between the diversification of the banana family and the geological and climatic history of southeast asia. New Phytologist, 210:1453–1465, 2016.

[66] J.D.G. Jones, B.J. Staskawicz, and J.L. Dangl. The plant immune system: From discovery to deployment. Cell, 187(9):2095–2116, 2024.

[67] L.M. Jubic, S. Saile, O.J. Furzer, F. El Kasmi, and J.L. Dangl. Help wanted: helper nlrs and plant immune responses. Current Opinion in Plant Biology, 50:82–94, 2019.

[68] J. Jumper, R. Evans, A. Pritzel, T. Green, M. Figurnov, O. Ronneberger, et al. Highly accurate protein structure prediction with alphafold. Nature, 596(7873):583–589, 2021.

[69] F. Jupe, L. Pritchard, G.J. Etherington, K. Mackenzie, P.J. Cock, F. Wright, et al. Identification and localisation of the nb-lrr gene family within the potato genome. BMC Genomics, 13(1):75, 2012.

[70] J. Jurka, V.V. Kapitonov, A. Pavlicek, P. Klonowski, O. Kohany, and J. Walichiewicz. Repbase update, a database of eukaryotic repetitive elements. Cytogenetic and Genome Research, 110(1-4):462–467, 2005.

[71] I. Kalvari, E.P. Nawrocki, N. Ontiveros-Palacios, J. Argasinska, K. Lamkiewicz, M. Marz, et al. Rfam 14: expanded coverage of metagenomic, viral and microrna families. Nucleic Acids Research, 49(D1):D192–D200, 2021.

[72] A. Kessel and N. Ben-Tal. Introduction to proteins: Structure, function, and motion. Chapman and Hall/CRC, 2nd edition, 2018.

[73] K. Khalturin, G. Hemmrich, S. Fraune, R. Augustin, and T.C. Bosch. More than just orphans: are taxonomically-restricted genes important in evolution? Trends in Genetics, 25(9):404–413, 2009.

[74] S.A. Kidane, B.H. Meressa, S. Haukeland, T. Hvoslef-Eide, C. Magnusson, M. Couvreur, et al. Occurrence of plant-parasitic nematodes on enset (*Ensete ventricosum*) in ethiopia with focus on pratylenchus goodeyi as a key species of the crop. Nematology, 23(5):529–541, 2020.

[75] D. Kim, J.M. Paggi, C. Park, C. Bennett, and S.L. Salzberg. Graph-based genome alignment and genotyping with hisat2 and hisat-genotype. Nature Biotechnology, 37(8):907–915, 2019.

[76] W. Kim, M. Mirdita, E.L. Karin, C.L.M. Gilchrist, H. Schweke, J. Söding, et al. Rapid and sensitive protein complex alignment with foldseek-multimer. Cold Spring Harbor Laboratory, 2024.

[77] Y. Kodama, M. Shumway, and R. Leinonen. The sequence read archive: explosive growth of sequencing data. Nucleic Acids Research, 40(D1):D54–D56, 2012.

[78] I. Korf. Gene finding in novel genomes. BMC Bioinformatics, 5, 2004.

[79] J. Kourelis and H. Adachi. Activation and regulation of nlr immune receptor networks. Plant and Cell Physiology, 63:1366–1377, 2022.

[80] J. Kourelis, T. Sakai, H. Adachi, and S. Kamoun. Refplantnlr is a comprehensive collection of experimentally validated plant disease resistance proteins from the nlr family. PLoS Biology, 19(10):e3001124, 2021.

[81] I. Letunic and P. Bork. Interactive tree of life (itol) v5: an online tool for phylogenetic tree display and annotation. Nucleic Acids Research, 49(W1):W293–W296, 2021.

[82] H. Li. Minimap2: pairwise alignment for nucleotide sequences. Bioinformatics, 34:3094–3100, 2018.

[83] W. Li and A. Godzik. Cd-hit: a fast program for clustering and comparing large sets of protein or nucleotide sequences. Bioinformatics, 22:1658–1659, 2006.

[84] X. Liu, R. Arshad, X. Wang, W.-M. Li, Y. Zhou, X.-J. Ge, et al. The phased telomere-to-telomere reference genome of musa acuminata, a main contributor to banana cultivars. Scientific Data, 10(1), 2023.

[85] Y. Liu, Z. Zeng, Y.M. Zhang, Q. Li, X.M. Jiang, Z. Jiang, et al. An angiosperm nlr atlas reveals that nlr gene reduction is associated with ecological specialization and signal transduction component deletion. Molecular Plant, 14(12):2015–2031, 2021.

[86] J. Logemann, J. Schell, and L. Willmitzer. Improved method for the isolation of rna from plant tissues. Analytical Biochemistry, 163(1):16–20, 1987.

[87] A. Lomsadze, V. Ter-hovhannisyan, Y.O. Chernoff, and M. Borodovsky. Gene identification in novel eukaryotic genomes by self-training algorithm. Nucleic Acids Research, 33:6494–6506, 2005.

[88] L. Lu, J. Chen, S.M.C. Robb, Y. Okumoto, J.E. Stajich, and S.R. Wessler. Tracking the genome-wide outcomes of a transposable element burst over decades of amplification. Proceedings of the National Academy of Sciences USA, 114(49):E10550–E10559, 2017.

[89] M. Manni, M.R. Berkeley, M. Seppey, F.A. Simão, and E.M. Zdobnov. Busco update: Novel and streamlined workflows along with broader and deeper phylogenetic coverage for scoring of eukaryotic, prokaryotic, and viral genomes. Molecular Biology and Evolution, 38(10):4647–4654, 2021.

[90] G. Marçais, A.L. Delcher, A.M. Phillippy, R. Coston, S.L. Salzberg, and A. Zimin. Mummer4: A fast and versatile genome alignment system. PLoS Computational Biology, 14(1):e1005944, 2018.

[91] G. Marçais and C. Kingsford. A fast, lock-free approach for efficient parallel counting of occurrences of k-mers. Bioinformatics, 27(6):764–770, 2011.

[92] M. Martin. Cutadapt removes adapter sequences from high-throughput sequencing reads. EMBnet.journal, 17(1):10, 2011.

[93] N. Maruta, H. Burdett, B.Y.J. Lim, X. Hu, S. Desa, M.K. Manik, et al. Structural basis of nlr activation and innate immune signalling in plants. Immunogenetics, 74(1):5–26, 2022.

[94] B. Meszaros, P. Tompa, I. Simon, and Z. Dosztanyi. Molecular principles of the interactions of disordered proteins. Journal of Molecular Biology, 372(2):549–561, 2007.

[95] C. Mhiri, J.B. Morel, S. Vernhettes, J.M. Casacuberta, H. Lucas, and M.A. Grandbastien. The promoter of the tobacco tnt1 retrotransposon is induced by wounding and by abiotic stress. Plant Molecular Biology, 33(2):257–266, 1997.

[96] L.S. Mofatto, F.A. Carneiro, N.G. Vieira, K.E. Duarte, R.O. Vidal, J.C. Alekcevetch, et al. Identification of candidate genes for drought tolerance in coffee by high-throughput sequencing in the shoot apex of different coffea arabica cultivars. BMC Plant Biology, 16:94, 2016.

[97] H. Moon, A.R. Jeong, O.K. Kwon, and C.J. Park. Oryza-specific orphan protein triggers enhanced resistance to xanthomonas oryzae pv. oryzae in rice. Frontiers in Plant Science, 13:859375, 2022.

[98] N. Murvai, L. Kalmar, B. Szabo, E. Schad, A. Micsonai, J. Kardos, et al. Cellular chaperone function of intrinsically disordered dehydrin erd14. International Journal of Molecular Sciences, 22(12), 2021.

[99] S. Muzemil. Manuscript genome annotation and analysis files. Zenodo, 2026.

[100] S. Muzemil, A. Chala, B. Tesfaye, D.J. Studholme, M. Grant, Z. Yemataw, et al. Evaluation of 20 enset (*Ensete ventricosum*) landraces for response to *Xanthomonas vasicola* pv. *Musacearum* infection. European Journal of Plant Pathology, 161(4):821–836, 2021.

[101] S. Muzemil, T.M. Olango, and Z. Yemataw. Enset (*Ensete ventricosum*) landrace database (version 0), 2023.

[102] G.V. Nakato, P. Christelová, E. Were, M. Nyine, T.A. Coutinho, J. Doležel, et al. Sources of resistance in musa to *Xanthomonas campestris* pv. *musacearum*, the causal agent of banana xanthomonas wilt. Plant Pathology, 68, 2019.

[103] V. Nakato, G. Mahuku, and T. Coutinho. Xanthomonas campestris pv. musacearum: a major constraint to banana, plantain and enset production in central and east africa over the past decade. Molecular Plant Pathology, 19(3):525–536, 2018.

[104] E.P. Nawrocki and S.R. Eddy. Infernal 1.1: 100-fold faster rna homology searches. Bioinformatics, 29:2933–2935, 2013.

[105] S. Ou and N. Jiang. Ltr_finder_parallel: Parallelization of ltr_finder enabling rapid identification of long terminal repeat retrotransposons. Mobile DNA, 10(40):1–3, 2019.

[106] S. Ou, W. Su, Y. Liao, K. Chougule, J.R.A. Agda, A.J. Hellinga, et al. Benchmarking transposable element annotation methods for creation of a streamlined, comprehensive pipeline. Genome Biology, 20:275, 2019.

[107] A.B. Pereira, M. Marano, R. Bathala, R.A. Zaragoza, A. Neira, A. Samano, et al. Orphan genes are not a distinct biological entity. Bioessays, 47(1):e2400146, 2025.

[108] L.T.J. Pijls, A.A.M. Timmer, Z. Wolde-Gebriel, and C.E. West. Cultivation, preparation and consumption of ensete (*Ensete ventricosum*) in ethiopia. Journal of the Science of Food and Agriculture, pages 1–11, 1995.

[109] A.R. Quinlan and I.M. Hall. Bedtools: a flexible suite of utilities for comparing genomic features. Bioinformatics, 26(6):841–842, 2010.

[110] E. Ramallo, R. Kalendar, A.H. Schulman, and J.A. Martínez-Izquierdo. Reme1, a copia retrotransposon in melon, is transcriptionally induced by uv light. Plant Molecular Biology, 66(1-2):137–150, 2008.

[111] Fidel Ramírez, Friederike Dündar, Sarah Diehl, Björn A. Grüning, and Thomas Manke. deeptools: a flexible platform for exploring deep-sequencing data. Nucleic Acids Research, 42(W1):W187–W191, 05 2014.

[112] S. Rana, P.R. Aggarwal, V. Shukla, U. Giri, S. Verma, and M. Muthamilarasan. Genome editing and designer crops for the future. In Methods in Molecular Biology, pages 37–69. Springer US, 2022.

[113] T.R. Ranallo-Benavidez, K.S. Jaron, and M.C. Schatz. Genomescope 2.0 and smudgeplot for reference-free profiling of polyploid genomes. Nature Communications, 11(1), 2020.

[114] M. Rocheta, L. Carvalho, W. Viegas, and L. Morais-Cecílio. Corky, a gypsy-like retrotransposon is differentially transcribed in quercus suber tissues. BMC Research Notes, 5(1):432, 2012.

[115] C. Sambles, L. Venkatesan, O.M. Shittu, J. Harrison, K. Moore, L. Tripathi, et al. Genome sequencing data for wild and cultivated bananas, plantains and abacá. Data in Brief, 33:106341, 2020.

[116] T. Sanderson, M.L. Bileschi, D. Belanger, and L.J. Colwell. Proteinfer, deep neural networks for protein functional inference. eLife, 12, 2023.

[117] E.W. Sayers, R. Agarwala, E.E. Bolton, J.R. Brister, K. Canese, K. Clark, et al. Database resources of the national center for biotechnology information. Nucleic Acids Research, 47(D1):D23–D28, 2019.

[118] E.W. Sayers, M. Cavanaugh, K. Clark, J. Ostell, K.D. Pruitt, and I. Karsch-Mizrachi. Genbank. Nucleic Acids Research, 47(D1):D94–D99, 2019.

[119] A. Schlessinger, C. Schaefer, E. Vicedo, M. Schmidberger, M. Punta, and B. Rost. Protein disorder—a breakthrough invention of evolution? Current Opinion in Structural Biology, 21(3):412–418, 2011.

[120] A.L. Schoonmaker, R.M. Hillabrand, V.J. Lieffers, P.S. Chow, and S.M. Landhäusser. Seasonal dynamics of non-structural carbon pools and their relationship to growth in two boreal conifer tree species. Tree Physiology, 41(9):1563–1582, 2021.

[121] J. Shi and C. Liang. Generic repeat finder: A high-sensitivity tool for genome-wide de novo repeat detection. Plant Physiology, 180:1803–1815, 2019.

[122] F. Sievers and D.G. Higgins. Clustal omega for making accurate alignments of many protein sequences. Protein Science, 27(1):135–145, 2018.

[123] Y. Song and J. Wang. ggcoverage: an r package to visualize and annotate genome coverage for various ngs data. BMC Bioinformatics, 24(1):309, 2023.

[124] A. Stamatakis. Raxml version 8: a tool for phylogenetic analysis and post-analysis of large phylogenies. Bioinformatics, 30(9):1312–1313, 2014.

[125] B. Steuernagel, K. Witek, S.G. Krattinger, R.H. Ramirez-Gonzalez, H.J. Schoonbeek, G. Yu, et al. The nlr-annotator tool enables annotation of the intracellular immune receptor repertoire. Plant Physiology, 183(2):468–482, 2020.

[126] D.J. Studholme, E. Wicker, S.M. Abrare, A. Aspin, A. Bogdanove, K. Broders, et al. Transfer of *Xanthomonas campestris* pv. *arecae* and *X. campestris* pv. *musacearum* to *X. vasicola* (vauterin) as *X. vasicola* pv. *arecae* comb. nov. and *X. vasicola* pv. *musacearum* comb. nov. Phytopathology, 110(6):1153–1160, 2020.

[127] W. Su, X. Gu, and T. Peterson. Tir-learner, a new ensemble method for tir transposable element annotation, provides evidence for abundant new transposable elements in the maize genome. Molecular Plant, 12:447–460, 2019.

[128] H. Sun, J. Ding, M. Piednoël, and K. Schneeberger. findgse: estimating genome size variation within human and arabidopsis using k-mer frequencies. Bioinformatics, 34(4):550–557, 2018.

[129] M. Tarailo-Graovac and N. Chen. Using repeatmasker to identify repetitive elements in genomic sequences. Current Protocols in Bioinformatics, 2009.

[130] D.E. Tarr and H.M. Alexander. Tir-nbs-lrr genes are rare in monocots: evidence from diverse monocot orders. BMC Research Notes, 2:197, 2009.

[131] R.L. Tatusov, E.V. Koonin, and D.J. Lipman. A genomic perspective on protein families. Science, 278(5338):631–637, 1997.

[132] D. Tautz and T. Domazet-Lošo. The evolutionary origin of orphan genes. Nature Reviews Genetics, 12(10):692–702, 2011.

[133] H. Thorvaldsdóttir, J.T. Robinson, and J.P. Mesirov. Integrative genomics viewer (igv): high-performance genomics data visualization and exploration. Briefings in Bioinformatics, 14(2):178–192, 2013.

[134] P. Tompa and D. Kovacs. Intrinsically disordered chaperones in plants and animals. Biochemistry and Cell Biology, 88(2):167–174, 2010.

[135] R. Trivedi and H.A. Nagarajaram. Intrinsically disordered proteins: An overview. International Journal of Molecular Sciences, 23(22), 2022.

[136] A. Tsegaye and P.C. Struik. Enset (*Ensete ventricosum* (welw.) cheesman) kocho yield under different crop establishment methods as compared to yields of other carbohydrate-rich food crops. NJAS: Wageningen Journal of Life Sciences, 49(1):81–94, 2001.

[137] M. Van Kempen, S.S. Kim, C. Tumescheit, M. Mirdita, J. Lee, C.L.M. Gilchrist, et al. Fast and accurate protein structure search with foldseek. Nature Biotechnology, 42(2):243–246, 2024.

[138] M. Vasimuddin, S. Misra, H. Li, and S. Aluru. Efficient architecture-aware acceleration of bwa-mem for multicore systems. In 2019 IEEE International Parallel and Distributed Processing Symposium (IPDPS), pages 314–324, 2019.

[139] C. Vitte, M.A. Fustier, K. Alix, and M.I. Tenaillon. The bright side of transposons in crop evolution. Briefings in Functional Genomics, 13(4):276–295, 2014.

[140] Z. Wang, H. Miao, J. Liu, B. Xu, X. Yao, C. Xu, et al. *Musa balbisiana* genome reveals subgenome evolution and functional divergence. Nature Plants, 5:810–821, 2019.

[141] N.I. Weisenfeld, V. Kumar, P. Shah, D.M. Church, and D.B. Jaffe. Direct determination of diploid genome sequences. Genome Research, 27(5):757–767, 2017.

[142] T. Wicker, F. Sabot, A. Hua-Van, J.L. Bennetzen, P. Capy, B. Chalhoub, et al. A unified classification system for eukaryotic transposable elements. Nature Reviews Genetics, 8:973–982, 2007.

[143] H. Wickham, M. Averick, J. Bryan, W. Chang, L. McGowan, R. François, et al. Welcome to the tidyverse. Journal of Open Source Software, 4(43):1686, 2019.

[144] P. Woodrow, G. Pontecorvo, L.F. Ciarmiello, A. Fuggi, and P. Carillo. Ttd1a promoter is involved in dna-protein binding by salt and light stresses. Molecular Biology Reports, 38(6):3787–3794, 2011.

[145] World Bank. Databank: Population estimates and projections filter by sub-saharan africa (excluding high income) and using time 2025. https://databank.worldbank.org/source/population-estimates-and-projections, 2025.

[146] W. Xiao, H. Liu, Y. Li, X. Li, C. Xu, M. Long, et al. A rice gene of de novo origin negatively regulates pathogen-induced defense response. PLoS One, 4(2):e4603, 2009.

[147] W. Xiong, L. He, J. Lai, H.K. Dooner, and C. Du. Helitronscanner uncovers a large overlooked cache of helitron transposons in many plant genomes. Proceedings of the National Academy of Sciences USA, 111(28):10263–10268, 2014.

[148] Z. Yemataw, G. Blomme, S. Muzemil, and K. Tesfaye. Assessing qualitative and phenotypic trait diversity in ethiopian enset (*Ensete ventricosum* (welw.) cheesman) landraces. Fruits, 73(6):310–327, 2018.

[149] Z. Yemataw, A. Chala, D. Ambachew, D. Studholme, M. Grant, and K. Tesfaye. Morpho-logical variation and inter-relationships of quantitative traits in enset (ensete ventricosum (welw.) cheesman) germplasm from south and south-western ethiopia. Plants, 6(4):56, 2017.

[150] Z. Yemataw, H. Mohamed, M. Diro, T. Addis, and G. Blomme. Ethnic-based diversity and distribution of enset (*Ensete ventricosum*) clones in southern ethiopia. Journal of Ecology and The Natural Environment, 6:244–251, 2014.

[151] Z. Yemataw, S. Muzemil, D. Ambachew, L. Tripathi, K. Tesfaye, A. Chala, et al. Genome sequence data from 17 accessions of *Ensete ventricosum*, a staple food crop for millions in ethiopia. Data in Brief, 18:285–293, 2018.

[152] Z. Yemataw, K. Tesfaye, M. Grant, D.J. Studholme, and A. Chala. Multivariate analysis of morphological variation in enset (*Ensete ventricosum* (welw.) cheesman) reveals regional and clinal variation in germplasm from south and south western ethiopia. Australian Journal of Crop Science, 12(12):1849–1858, 2018.

[153] Z. Yemataw, K. Tesfaye, A. Zeberga, and G. Blomme. Exploiting indigenous knowledge of subsistence farmers for the management and conservation of enset (*Ensete ventricosum* (welw.) cheesman) (musaceae family) diversity on-farm. Journal of Ethnobiology and Ethnomedicine, 12(1):34, 2016.

[154] D. Yirgou and J.F. Bradbury. Bacterial wilt of enset (*Ensete ventricosum*) incited by *Xanthomonas musacearum* sp.n. Phytopathology, 58(1):111–112, 1968.

[155] D. Yirgou and J.F. Bradbury. A note on wilt of banana caused by the enset wilt organism *Xanthomonas musacearum*. East African Agricultural and Forestry Journal, 14:111–114, 1974.

[156] A. Yu, G. Lepère, F. Jay, J. Wang, L. Bapaume, Y. Wang, et al. Dynamics and biological relevance of dna demethylation in arabidopsis antibacterial defense. Proceedings of the National Academy of Sciences USA, 110(6):2389–2394, 2013.

[157] R.G. Zhang, G.Y. Li, X.L. Wang, J. Dainat, Z.X. Wang, S. Ou, et al. Tesorter: an accurate and fast method to classify ltr-retrotransposons in plant genomes. Horticulture Research, 9:uhac017, 2022.

